# In mice, a population of male germ cells show characteristics of non-apoptotic cell death during G0 arrest

**DOI:** 10.64898/2026.05.07.723530

**Authors:** Kara Stark, Talia Hatkevich, Edward Miao, Tomokazu Souma, Blanche Capel

**Affiliations:** Department of Cell Biology, Duke University School of Medicine, Durham, NC, USA; Department of Integrative Immunobiology, Duke University School of Medicine, Durham, NC, USA; Department of Molecular Genetics and Microbiology, Duke University School of Medicine, Durham, NC, USA; Department of Pathology, Duke University School of Medicine, Durham, NC, USA; Division of Nephrology, Department of Medicine, Duke University School of Medicine, Durham, NC, USA

**Keywords:** Germ cells, Cell death, G0 arrest

## Abstract

In mammals, a small population of spermatogonial stem cells (SSCs) is established shortly after birth. These cells self-renew and produce sperm for the entirety of a male’s reproductive lifespan, passing the genome on to the next generation. Thus, establishment of a population of SSCs with high genomic integrity is essential. SSCs are derived from a much larger precursor population of male germ cells (MGCs) that differentiate during fetal life. During the last third of gestation, MGCs undergo a prolonged period of G0 cell cycle arrest during which they sustain high levels of transcription and acquire epigenetic programming for SSC fate. Although these differentiation steps can cause cellular and genomic damage, it has been unclear whether selection for germ cell quality occurs during G0 arrest since no classic markers of cell death have been detected. In this study, we utilize a mouse model to characterize a population of MGCs that begin to accumulate markers if cell death, such as AnnexinV (AnV) and propidium iodide (PI), at E16.5. The AnV- and PI-positive MGC population is characterized by low expression of the RNA-binding protein, Dead End 1 (DND1), and exhibit dsDNA breaks and mitochondrial dysfunction. Interestingly, we do not see evidence of an active cell death cascade until the time of birth, where we see phosphorylation of MLKL, a hallmark of a necroptotic cell death mechanism. Based on these findings, we propose that variable cellular health is an important basis for selection of the SSC precursors.

**Significance Statement:** Spermatogonial stem cells (SSCs) are essential for reproductive fitness, yet how their precursors are selected during development is not known. Utilizing a mouse model, this study describes high levels of cellular damage within a subset of male germ cells (MGCs) during G0 arrest. The damaged MGC population was marked by low expression of the RNA-binding protein, DND1, and was strongly associated with mitochondrial dysfunction and dsDNA breaks. We observed signs of non-apoptotic cell death by embryonic day (E)16.5 and the appearance of necroptotic markers in MGCs at the time of birth. This study uncovers previously unknown heterogeneity in the MGC pool and points to MGC health as an important source of selection during G0 arrest.

## Introduction

Spermatogonial stem cells (SSCs) are the resident stem cells of the testis and are responsible for both self-renewing and producing sperm for the entirety of a male’s reproductive lifespan. SSCs are derived from primordial germ cells (PGCs), which are specified early in embryonic development. After their specification, PGCs undergo genome-wide DNA demethylation as they migrate through the gut mesentery to the developing gonad[1]. Once inside the gonad, PGCs receive signals from the surrounding soma that initiate the acquisition of sex-specific characteristics in a process known as germ cell sex determination. In the testis, male germ cells (MGCs) undergo several rounds of mitotic divisions before entering a prolonged period of G0 cell cycle arrest lasting, in mice, from embryonic day (E) 14.5 until postnatal day (P) 1. G0 arrest is a period of intense epigenetic remodeling that prepares the multipotent MGCs to become unipotent prospermatogonia [2, 3]. After G0 exit, a select population of prospermatogonia, migrate to the basement membrane of the testis cords and establish the SSC population.

SSCs contain the DNA that will be passed to the next generation; thus, the fidelity of the SSC genome is essential for evolutionary fitness. Despite the necessity for maintenance of genomic integrity, nearly every step of germ cell development is conducive to DNA damage. Mechanical stress during PGC migration can induce DNA breaks, DNA demethylation can lead to ssDNA lesions, and DNA hypomethylation renders the genome vulnerable to transposon activation[4–6]. Additionally, epigenetic reprogramming can create DNA damage via the movement of histones and other proteins on and off the DNA, and G0-stage MGCs experience hypertranscriptional activity which can induce R-loop formation and DNA breaks[7–9]. Thus, each step of SSC development imposes many threats to the germ cell genome.

Interestingly, despite their hazardous developmental path, SSCs maintain a very low mutational load compared to somatic cells[10–13]. After G0 exit, the majority of MGCs begin to differentiate and enter the first wave of spermatogenesis, and only a small proportion of MGCs migrate to the basement membranes and take up SSC fate[14]. Previous studies have found that the MGCs that become SSCs also have a lower mutational load than those that enter the first wave, and both these germ cell populations have lower mutational rates than somatic cells[10, 11]. This suggests that there are stringent quality control mechanisms in place that safeguard MGC development and govern SSC selection.

Developmental quality control mechanisms may function either through the implementation of (1) unique cellular mechanisms to mitigate damage, or (2) efficient strategies for the removal of damaged cells. There is evidence that MGCs utilize both these approaches to ensure SSC genomic fidelity. For example, the piRNA pathway is an MGC-specific strategy for transposon silencing that combats selfish genetic elements during long periods of epigenetic vulnerability[15, 16]. Additionally, two well-characterized waves of apoptosis occur during MGC development, at E13.5 and P10[17–21]. MGC apoptosis at E13.5 functions to remove damaged or improperly programmed MGCs from the testis prior to G0 arrest[18, 20], whereas apoptosis within the first month of life modulates the number of germ cells that can be supported by the somatic cells of the testes[19, 21]. While less work has been done to characterize cellular health during G0 arrest, Wang *et al.* reported in 1998 the appearance of necrotic MGCs in EM images of testis cords during late gestation and early in the neonatal period[17]. However, to our knowledge, no one has followed up on this observation, and thus variation in MGC health during G0 arrest has gone unexplored.

Using a GFP reporter line, we previously identified differential expression of the germ cell-specific RNA-binding protein, Dead End 1 (DND1), within the MGC population[22]. Beginning at E16.5, DND1-hi and DND1-lo cell populations could be distinguished through FACS or transcriptome analysis[23]. Additionally, we previously showed that the DND1-lo cells are positive for the early cell death marker AnnexinV (AnV). However, this result was confusing as markers of apoptosis are absent during this developmental window[17, 23]. Additionally, while DND1 is known to be a necessary factor for MGC development, the functional significance of the DND1-hi and -lo cell populations remained unclear. In this study, we describe a population of MGCs that show molecular hallmarks of necroptotic cell death at the time of birth. During mid- to -late gestation, we found that ∼70% of MGCs are positive for propidium iodide (PI). This population, which is marked by low DND1 expression, also displays an increase in dsDNA breaks and exhibits signs of mitochondrial dysfunction. Notably, we found that mitochondrial stress could prematurely induce DND1-lo expression at E14.5, leading us to propose that differential metabolic health is a major determinant of prospermatogonial selection during G0 arrest.

## Results

### G0-staged MGCs show signs of non-apoptotic cell death

A previous study from our lab reported that, at E16.5, a subpopulation of MGCs become positive for the cell death marker, AnV[23]. This result was surprising because AnV is typically considered an early apoptotic marker, and no MGC apoptosis has been previously reported within this period[17]. To assess MGC health during G0 arrest, we performed flow analysis of G0-staged testes with AnV and propidium iodide (PI). Cell stress can cause phosphatidylserine residues to be flipped to the outer leaflet of the cell membrane, which is detected with AnV. If a cell recovers from a stressor, plasma membrane asymmetry can be restored, leading to loss of the AnV signal[24]. Alternatively, cells undergoing diverse mechanisms of cell death become positive for both AnV and PI, a DNA stain that can only penetrate the nucleus after plasma membrane permeabilization[25, 26]. To assess MGC health during G0 arrest, we performed flow analysis on G0-staged testes with AnV and PI. Utilizing DND1-GFP expression[22] to isolate the MGC population via flow cytometry (Fig. 1A), we measured the number of AnV and PI positive cells and observed a significant increase between E14.5 and E16.5. (Fig. 1B, C). The presence of AnV and PI together are considered indicative of active cell death[25, 26]; thus, these data support the presence of an actively dying MGC population at E16.5.

**Figure 1.**
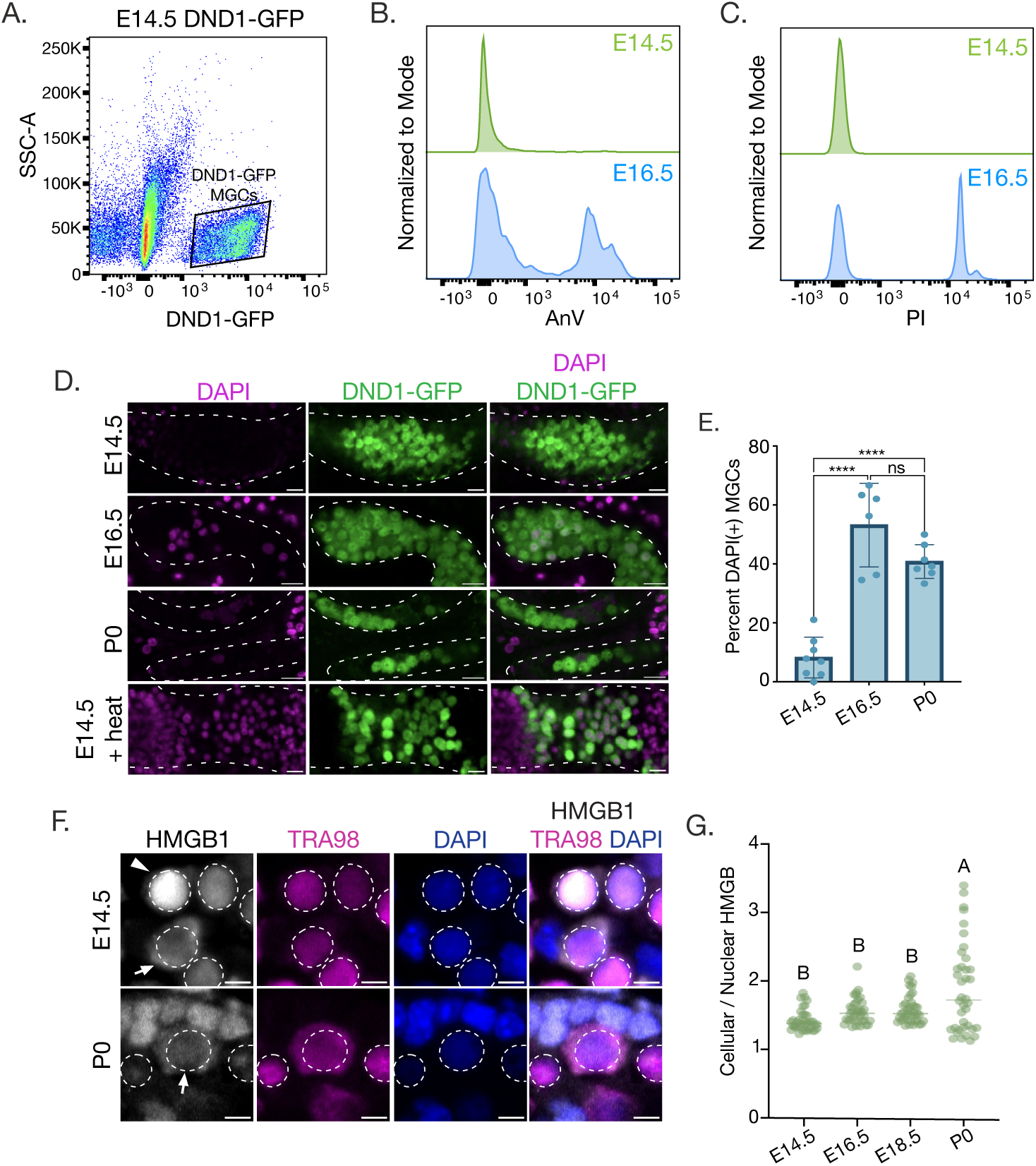
Some MGCs show signs of non-apoptotic cell death during G0 arrest. ***(A)*** Scatter plot demonstrating flow cytometry isolation of DND1*-*GFP expressing MGC population at E14.5. ***(B, C)*** Geometric mean of AnV *(B)* or PI *(C)* signal in E14.5 (green) and E16.5 (blue) MG*C*s. ***(D)*** Representative images of live DND1*-*GFP testis cords, DAPI used to label damaged cells; testis cords outlined with dashed line, scale bar = 20µm. ***(E)*** Bar graph representing the number of DAPI-positive MGCs per testis cord; individual points represent the percentage of DAPI-positive MGCs per image analyzed. Biological replicates n = 2 per stage. Statistical analysis based on ANOVA and Tukey’s multiple comparison’s test. ***(F)*** Representative images of HMGB1 sub-cellular localization at E14.5 and P0. Panels display HMGB1 (grey), TRA98 (magenta), and DAPI (blue). MGC nuclei outlined with dashed line; triangle indicates MGC with nuclear HMGB1 localization, arrow*s* indicate cell*s* with cytoplasmic HMGB1. Scale bar = 5µm. ***(G)*** Quantification of the sum fluorescent intensity of cellular HMGB1 signal over nuclear HMGB1 signal across G0 arrest. Statistical analysis based on ANOVA and Tukey’s multiple comparison’s test. Analysis showed that the pixel intensity, or mean grey value (MGV), at P0 (n = 38 cells) was statistically different than all other stages tested (A, *P* < 0.001). There was no statistical difference (B, *P* > 0.05) in between E14.5 (n = 38 cells), E16.5, (n = 37 cells) and E18.5 (n = 40 cells).

To confirm that the observed MGC damaged at E16.5 was not an artifact of testis dissociation, we performed live-imaging of DND1-GFP expressing testes incubated with low concentrations of DAPI. Although DAPI can be a semi-permeable marker in living tissue, when used at low concentrations, DAPI only accumulates in the nuclei of damaged cells[27, 28]. The benefit of this approach is that it does not require tissue dissociation to examine nuclear permeability, decreasing the risk of experimentally induced damage. At E14.5, most MGCs were negative for DAPI signal (91.77±6.88%, Fig. 1D, E), consistent with the finding that E14.5 MGCs are negative for AnV/PI (Fig. 1B, C). However, incubation of E14.5 testes at 65°C prior to DAPI incubation induced a large DAPI-positive population (∼100%, Fig. 1D), validating that DAPI can permeate the testis cords and label the nuclei of damaged cells. At E16.5 and P0, live imaging of testis cords revealed an overlap between DND1-GFP and DAPI signal (53.19±14.19% and 40.82±5.72% at E16.5 and P0, respectively, Fig. D, E). These data support the presence of damaged MGCs within intact testis cords during G0 arrest.

During development, two waves of MGC apoptosis have been characterized at E13.5 and P10[17–21]. However, both EM imaging and TUNEL staining have confirmed the absence of apoptotic MGCs during G0 arrest[17]. We stained for the canonical apoptotic marker, cleaved-Caspase3 (cCASP3), and confirmed that G0-staged MGCs were negative for cCASP3 signal (Fig. S1A). To investigate signs of non-apoptotic cell death, we examined the sub-cellular localization of the DNA chaperone, High-Mobility Group Box-1 (HMGB1). Although HMGB1 remains nuclear during apoptotic cell death[29, 30], release of HMGB1 into the cytoplasm and extracellular space is associated with several non-apoptotic cell death pathways[29–31]. To quantify HMGB1 localization, we calculated the ratio of cellular to nuclear HMGB1; the measurement of cellular HMGB1 included signal from the nucleus, so a ratio of 1 would indicate 100% nuclear HMGB1. At E14.5, most MGC nuclei were brightly positive for HMGB1 and the MGC nuclear marker, TRA98, although a small proportion of MGCs did exhibit cytoplasmic HMGB1 (Fig. 1F, indicated with arrow). We found a similar pattern at E16.5 and E18.5, where the ratio of cellular to nuclear HMGB1 signal was <1.5 for most MGCs (Fig. 1G), indicating that the majority of HMGB1 protein is localized within the nucleus during late gestation. However, at P0, most MGCs exhibited a shift in HMGB1 localization into the cytoplasm, although HMGB1 remained nuclear in some MGCs. This was reflected by a wide range in the ratio of cellular to nuclear HMGB1, which was between 1 and 4 (Fig. 1F, G). Release of nuclear HMGB1 suggests the initiation of a non-apoptotic cell death mechanism at the time of birth.

During development, MGCs exist in clonally derived cysts. Both MGC clonality and their cystic nature play a role in germ cell survival and fate decisions[18, 32, 33]. In both *Drosophila* and in E13.5 male mice, male germ cells die during differentiation in a clonal manner[18, 33, 34]. To ask whether this is also true during G0 arrest, we crossed a male *Oct4-CreER* mouse to a female harboring a *Rosa-mT/mG* reporter. Pregnant females were injected with a low dose of tamoxifen at E11.5 to induce clonal MGC labeling with a membrane-targeted GFP (Fig. S1B). We then examined the localization pattern of HMGB1 within MGC clones and observed significant variation; individual clones consisted of both cells with high and low cellular/nuclear HMGB1 ratios (Fig. S1C, D). This suggests that either neonatal MGCs do not die as clones, or death is asynchronous among clonal members.

### Damaged MGCs show characteristics of necroptotic cell death at the time of birth

Next, we investigated whether MGCs showed characteristics of a specific form of cell death. A previous study reported the appearance of necrotic MGCs during G0 arrest[17]. Necroptosis is a programmed form of cell necrosis associated with the release of nuclear HMGB1[31, 35–37]. To test for necroptotic activation in the G0 staged mouse testis, we stained for phosphorylation of Mixed lineage kinase domain-like protein (MLKL), the terminal step of necroptosis. At embryonic stages of G0 arrest (E14.5, E16.5, and E18.5), there was very little pMLKL signal within the testis cords (Fig. S2A). However, at the time of birth, there was robust accumulation of pMLKL puncta, characteristic of MLKL phosphorylation and multimerization (Fig. 2A). Quantification of pMLKL mean fluorescent intensity (or mean grey value, MGV) confirmed a highly significant change in pMLKL levels between E18.5 and P0 (Fig. 4B). While the MGV was significantly lower at P2 compared to P0, we observed instances of pMLKL positive cells that exhibited nuclear pMLKL localization (Fig. 2A), which may indicate a secondary function of pMLKL in MGC breakdown[38–40].

**Figure 2.**
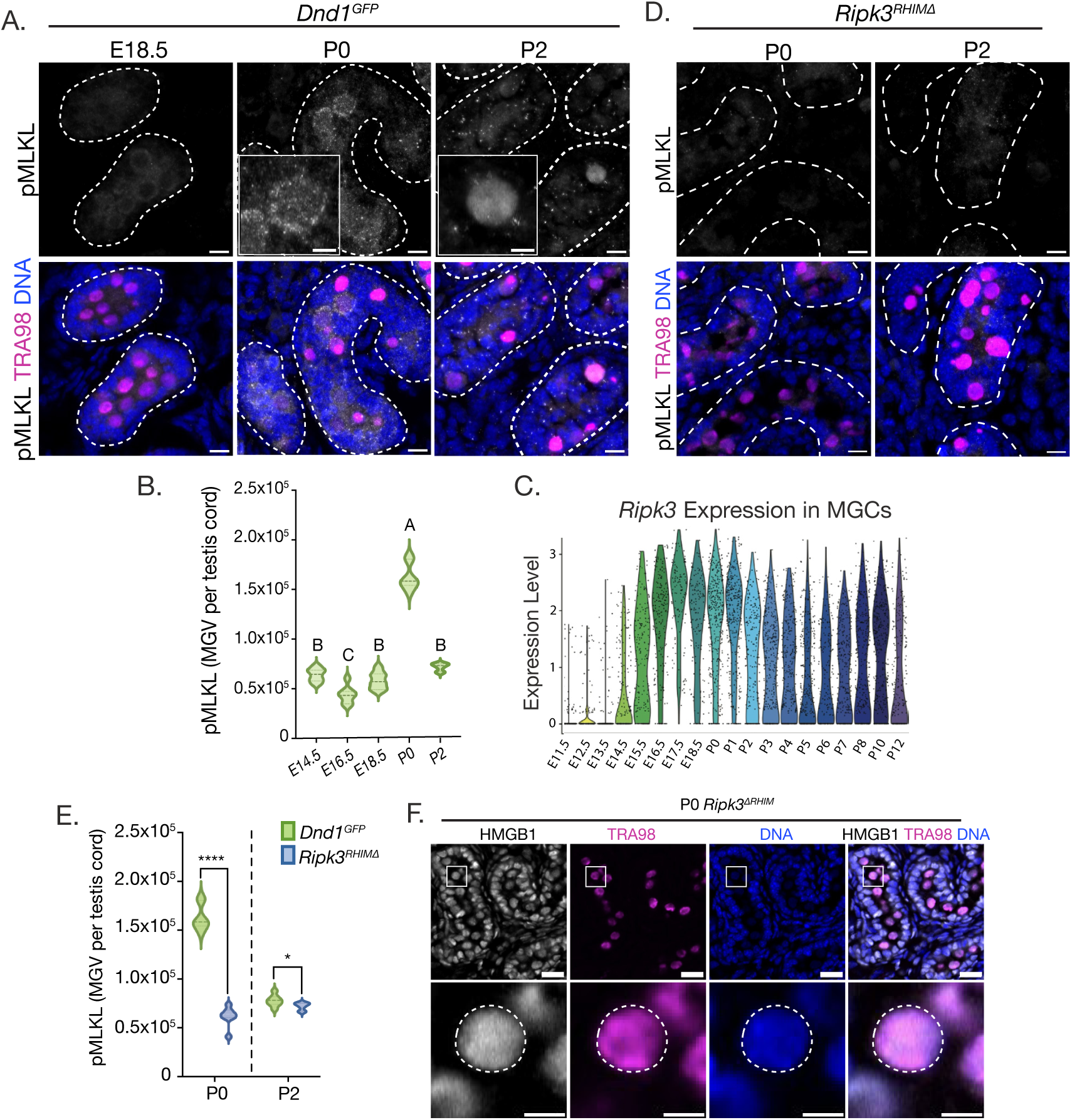
MGCs display hallmarks of necroptotic cell death at the time of birth. ***(A)*** Representative images of pMLKL levels at E18.5, P0, and P2. Images taken of sectioned testis tissue, testis cords outlined with dashed lines; bottom panels include DAPI and TRA98 staining. Scale bars = 10µm*. **(B)*** Quantification of pMLKL MGV per testis cord (n = 7 per stage). Statistical analysis based on ANOVA and Tukey’s multiple comparison’s test. MGV at P0 was statistically different than all other stages tested (A, *P* < 0.001). ***(C)*** Graph of *Ripk3* expression across germ cell development; data based on publicly available scRNA sequencing[43]. Expression level displayed on y-axis represents log(TPM/10 +1). ***(D)*** Representative images of pMLKL levels in the testes of *Ripk3^RHIMΔ^* mice. Images of sectioned testis tissue, testis cords outlined with dashed lines; bottom panels include DAPI and TRA98 staining. Scale bar = 10µm. ***(E)*** Quantification of pMLKL MGV per testis cord at P0 and P2 for *Dnd1^GFP^*and *Ripk3^RHIMΔ^* mice. Significance based on unpaired t test (n = 7 per stage, \**P* < 0.05, \*\*\*\**P* < 0.0001). ***(F)*** Representative images of HMGB1 sub-cellular localization at P0 in *Ripk3^RHIMΔ^* testes. Panels display HMGB1 (grey), TRA98 (magenta), and DAPI (blue). Nuclei outlined with dashed lines; Top panel scale bar = 10µm, bottom panel scale bar = 5µm.

During necroptosis, MLKL is often phosphorylated by a protein complex called the necrosome, which is mainly composed of Receptor-interacting serine/threonine-protein kinases 1 and 3 (RIPK1 and RIPK3) based on interactions between their RHIM domains[41, 42]. Interestingly, examination of publicly available scRNA sequencing data[43] revealed that expression of *Ripk3* was elevated specifically in G0-staged MGCs (Fig. 2C). To test whether MLKL phosphorylation depends on necrosome formation, we examined RIPK3 RHIM domain mutant testes and found a significant reduction in pMLKL signal at P0 and P2 (Fig. 2D,E). Previous literature has established that RIPK3 is necessary for release of nuclear HMGB1 during necroptosis[44]. In agreement with this, we found that *Ripk3^RHIMΔ^* MGCs maintained nuclear HMGB1 in the neonatal period, suggesting RIPK3 activity is necessary for the release of HMGB1 in neonatal MGCs (Fig. 2F). Necrosome formation can be downstream of Tumor necrosis factor (TNF) signaling[42], but knockout of TNF receptors 1 and 2 (*Tnfr1/2-KO*) did not significantly alter MKLK phosphorylation (Fig. S2B). We also tested whether Z-DNA sensing by Z-DNA binding protein 1 (ZBP1) contributes to MLKL phosphorylation and found that neonatal ZBP1 mutant testes exhibited no change in pMLKL levels (Fig. S2C).

Previous studies reported no significant change in MGC number between late gestation and the early neonatal period [45]. However, given the numerous markers of cell death MGCs display, we repeated this investigation. The levels of DND1-GFP decline sharply on P0, therefore we could not use this marker to sort or count MGCs. Therefore, we quantified MGC number by counting the number of TRA98-positive cells within 10µm tissue sections (Fig. S3A). Our results agreed with previous reports[45] showing that there is no change in MGC number spanning the fetal and neonatal periods. However, staining for the cell cycle marker Ki-67 revealed that a portion of the MGC population re-enters the cell cycle at P0 (Fig. S3B, C). Thus, we hypothesize that a decrease in MGC number between fetal and neonatal stages may be masked as some MGCs re-enter the cell cycle.

### MGCs that show signs of damage during GO arrest have low expression of DND1

Previous work from our lab established that the RNA-binding protein, DND1, is differentially expressed within the MGC population during G0 arrest, and that low expression of DND1 is associated with the cell death marker, AnV[23]. To determine whether other markers of cell damage characterize the DND1-lo cells we performed flow analysis of G0-staged cells from DND1-GFP testes stained with AnV and PI. In agreement with previous reports, E14.5 MGCs displayed a uniform distribution of DND1 expression (Fig. 3A), and 74.5% of MGCs were negative for both markers of cell death (Fig. 3C, C’). At E16.5, the MGC population becomes divided in their DND1-GFP signal, with roughly half becoming DND1-hi and half becoming DND1-lo (Fig. 3A’, B). Most DND1-hi cells (87.8%) were still negative for markers of cell death, but 73.1% of DND1-lo cells were positive for both AnV and PI (Fig. 3D, D’). Between E16.5 and E18.5, the DND1-lo population increased to ∼85% of the total MGC pool (Fig. 3A’’, B) and appeared to progress to late stages of cell death (63.6% were PI positive). However, 96.6% of the remaining DND1-hi cells were still negative for AnV and PI, suggesting these cells were still healthy at late stages of gestation (Fig. 3E, E’). These data indicate that DND1-lo marks a growing population of MGCs that progressively accumulate markers of cell damage during late gestation.

**Figure 3.**
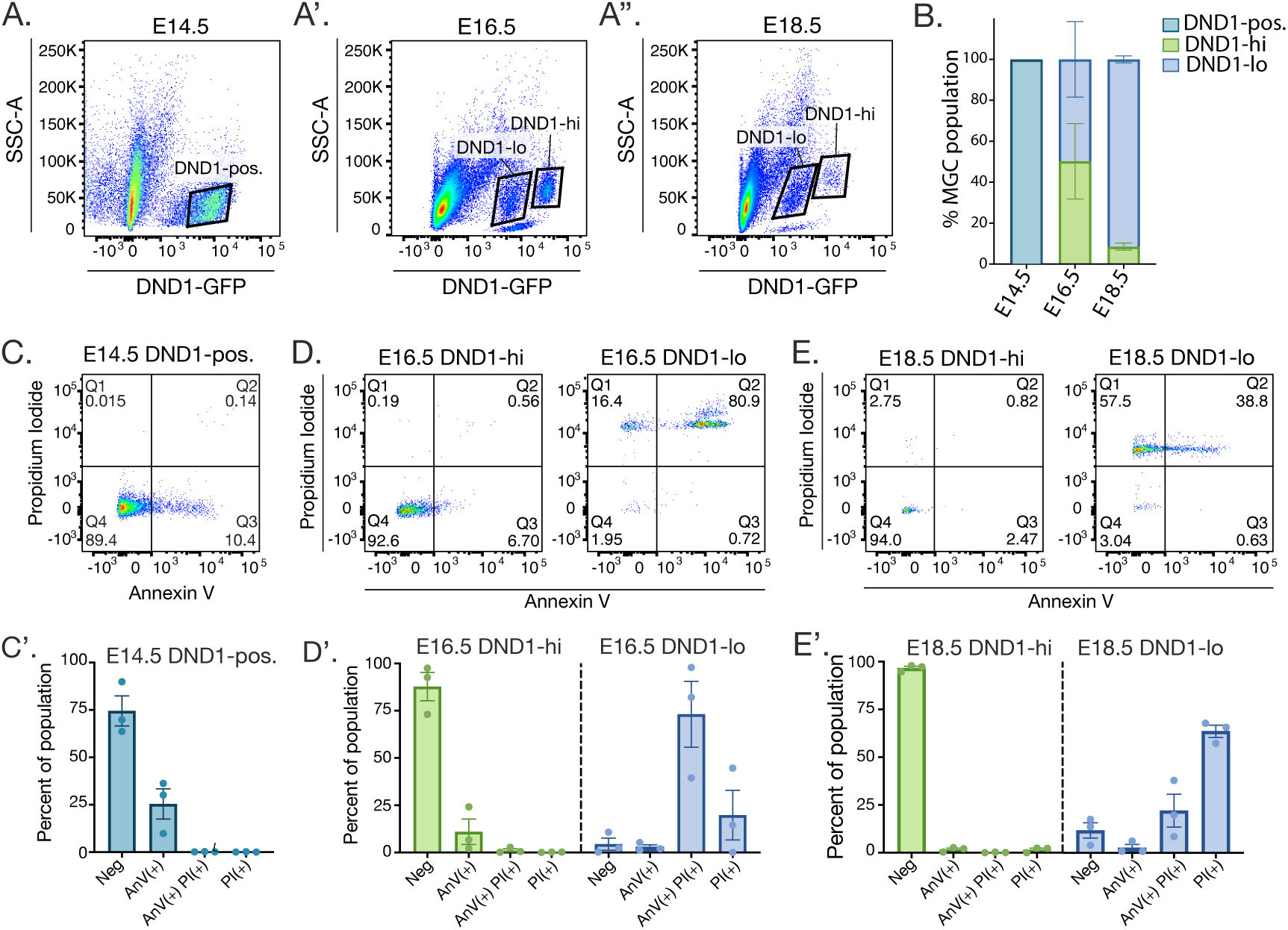
Cell death markers are associated with low expression of DND1. ***(A-A’’)*** Flow analysis of male gonadal cells from Dnd1^GFP/+^ mice at E14.5 *(A),* E16.5 *(A’),* and E18.5 *(A’’).* MGC population identified using DND1-GFP signal; DND1-hi and DND1-lo cells gated based off DND1-GFP signal intensity. ***(B)*** Graph of DND1-positive, DND1-hi, and DND1-lo MGC cells as a percentage of the total DND1-positive population at E14.5, E16.5 and E18.5. ***(C)*** Flow analysis of E14.5 DND1-positive MGCs stained with AnnexinV (AnV) and propidium iodide (PI). *(C’)* Graph indicating the percentage of MGCs negative for both AnV and PI, positive for AnV, positive for both AnV and PI, or positive for PI alone. ***(D, E)*** Flow analysis of DND1-hi (right) and DND1-lo (left) MGCs at E16.5 *(D)* and E18.5 *(E)* stained with AnV and PI. *(D’, E’)* Graphs indicating the percentage of DND1-hi and DND1-lo MGCs that are negative for both AnV and PI, positive for AnV, positive for both AnV and PI, or positive for PI alone at E16.5 *(D’)* and E18.5 *(E’).* All bar graphs represent the average percentages of a given MGC population (i.e. DND1-hi MGCs) over three litters (n = 3), error bars represent standard deviation.

Law *et al*. demonstrated that differential expression of the transcriptional repressor, ID4, during G0 arrest could identify the MGCs that will become SSCs (ID4-hi, “pre-SSC”) and the MGCs that will differentiate after birth and constitute the first wave of spermatogenesis (ID4-lo, “pre-first wave”). At E16.5, the ID4-hi population had higher expression of genes associated with SSC fate[46]. To ask whether there is a correlation between DND1 levels and fate determination, we examined previously published E16.5 DND1-hi and -lo bulk RNA sequencing data using the “pre-SSC’ and “pre-first wave” genes defined by Law *et al* (Fig. S4A). Unlike ID4, we found that DND1-hi cells had higher transcript levels of most genes examined, including those associated with a “pre-SSC” and “pre-first wave” path. This indicates that differential DND1 expression corresponds with differences in MGC health rather than SSC fate decisions. While most genes appeared higher in the DND1-hi population, we found that DND1-lo cells did have higher levels of some cell death response genes at E16.5, such as cytokines (*Cvcl5, Il11*), Cytokine receptors (*Ccr5, Cd40*), as well as some cell death regulatory genes (*Nfkb1, Ripk3, Cflar*, Fig. S4B). We hypothesize that these genes may be elevated in response to cell stress, as reflected by the accumulation of AnV/PI in the DND1-lo population.

### DND1-hi cells express higher levels of DNA repair proteins and contain lower instances of dsDNA breaks

To understand the differences between the DND1-hi and -lo populations that may contribute to a survival advantage of the DND1-hi MGCs, we performed proteomic analysis on DND1-hi and DND1-lo MGCs at E16.5. We identified a total of 5,737 proteins (Table S1). Interestingly, 18% of these proteins were differentially expressed between the DND1-hi and -lo populations, with 557 proteins significantly enriched in the DND1-hi cells and 497 proteins significantly enriched in the DND1-lo cells (Fig. 4A). GO term analysis of the differentially expressed proteins in DND1-hi cells showed enrichment of proteins involved in translation, protein biosynthesis, and metabolism (Fig. 4B). STRING analysis performed using enriched proteins from the DND1-hi population identified complexes involved in translation regulation, including many translation initiation factors and functional components of ribosomes. STRING analysis also identified enrichment of factors involved in transcriptional initiation. (Fig. 4C).

**Figure 4.**
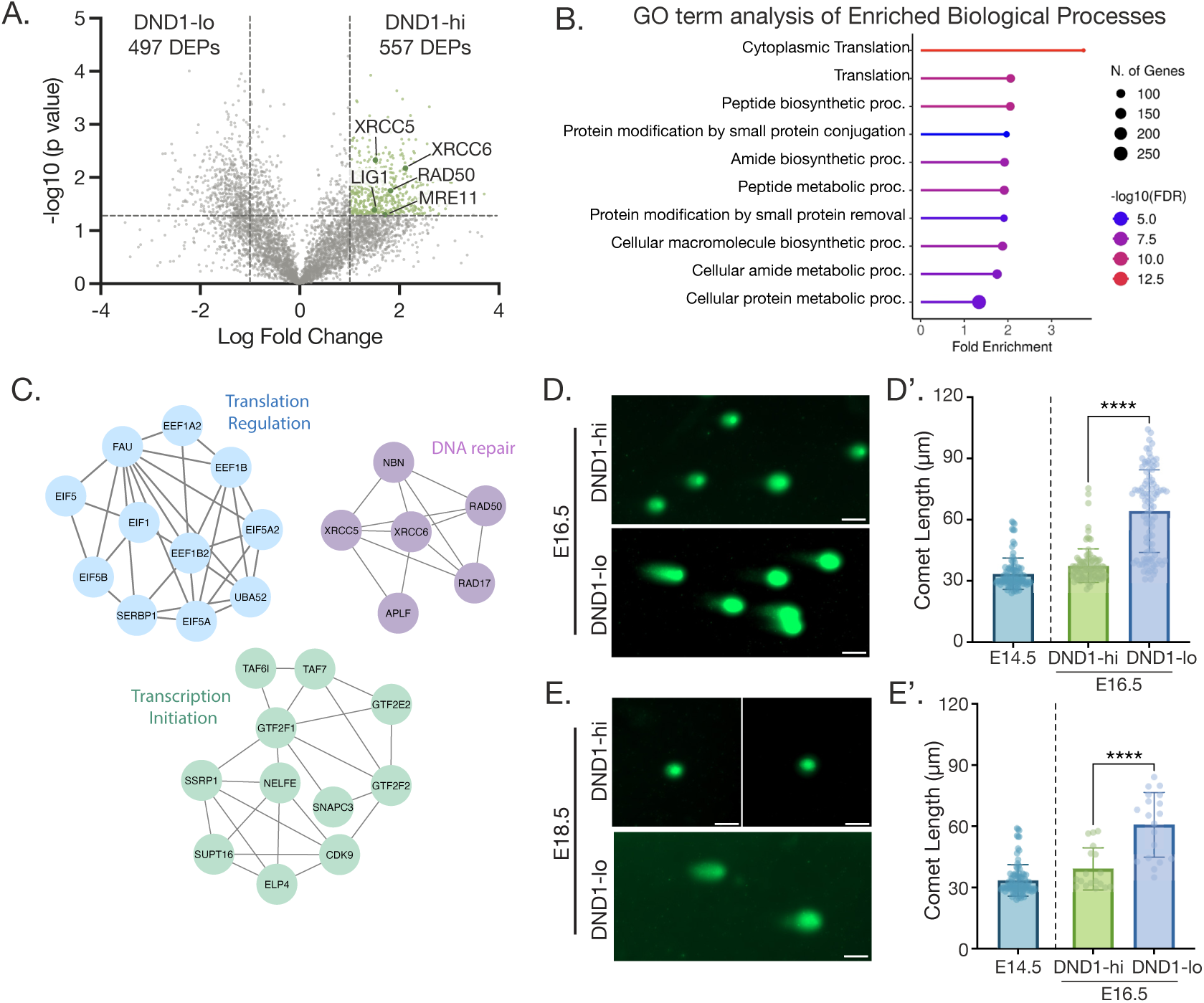
DND1-hi cells express higher levels of DNA repair proteins and translational machinery and have fewer dsDNA breaks. ***(A)*** Volcano plot comparing the proteomes of DND1-hi versus DND1-lo cell populations at E16.5. Differentially expressed proteins (DEPs) are defined as having a -log10(p-value) > 1.3 and a Log2FC < 1. DND1-hi DEPs are highlighted in green, and DNA repair machinery is labeled. ***(B)*** GO term analysis of DND1-hi DEPs. ***(C)*** STRING analysis performed on DND1-hi DEPs. Clusters represent proteins with known interactions. ***(D, E)*** Representative images of comet assay performed on flow sorted DND1-hi and DND1-lo cells at E16.5 *(D)* and E18.5 *(E).* Scale bar = 50µm. Quantification of E16.5 *(D’)* and E18.5 *(E’)* comet tail length in DND1-hi and -lo cells; E14.5 included on graphs for reference. Significance based on unpaired t test, *P* < 0.0001 (n = 100 nuclei for E16.5; n = 20 nuclei for E18.5).

Interestingly, STRING analysis of DND1-hi DEPs also identified enrichment of DNA-repair proteins including essential components of the MRN complex (MRE11, RAD50, and NBN) and other factors involved in non-homologous end-joining (Fig. 4A, C, S5A). Immunofluorescent staining detected high levels of *γ*H2AX at P2 but did not show abundant signal at E16.5 or E18.5 (Fig. S5B). This was surprising given the abundance of PI positive cells within the DND1-lo population at these stages (Fig. 3D, E). To ask whether DND1-lo cells harbor unmarked dsDNA breaks, possibly due to a lack of DNA repair machinery, we performed comet assays on flow sorted DND1-hi and DND1-lo cells. Notably, MGCs isolated via flow cytometry from E14.5 testes had very short comet tails, significantly shorter than those of somatic cells treated with heat (positive control) and more like those of untreated somatic cells (negative control, Fig. S5C, C’). Given < 0.3% of MGCs at E14.5 were positive for PI (Fig. 3C), this data is consistent with little DNA damage in E14.5 MGCs. However, at both E16.5 and E18.5, DND1-lo cells exhibited significantly longer comet tails than their stage-matched DND1-hi counterparts (Fig. 4D, D’). This suggests that, at E16.5 and E18.5, DND1-lo cells experience an increase in dsDNA breaks even though they are not positive for the classic marker, *γ*H2AX. Meanwhile, DND1-hi cells maintain high DNA fidelity, possibly due to their higher abundance of DNA repair proteins.

### DND1-lo cells show signs of mitochondrial dysfunction and have decreased levels of ROS

To further explore the state of DND1-lo cell health during G0 arrest, we examined the proteins that were enriched in the DND1-lo MGC proteome. GO term analysis of these proteins revealed significant enrichment for mitochondrial proteins, specifically proteins involved in electron transport chain (ETC) assembly (Fig. 5A, B). Cells undergoing mitochondrial dysfunction sense that their energetic needs are unmet and increase expression of ETC building blocks[47]; thus, we hypothesized that DND1-lo cells have low mitochondrial health. We performed flow analysis on dissociated gonads stained with TMRM, a cationic dye used to measure mitochondrial membrane potential. At E14.5, DND1-positive MGCs displayed uniform TMRM levels. However, E16.5 DND1-hi cells had significantly higher TMRM signal than the E16.5 DND1-lo or E14.5 MGCs (Fig. 5C, C’). This result was confirmed by repeating the experiment using MitoTracker, which also depends on positive membrane potential to label mitochondria. The DND1-hi cells exhibited a higher MitoTracker signal as compared to DND1-lo cells (Fig. S6A, B).

**Figure 5.**
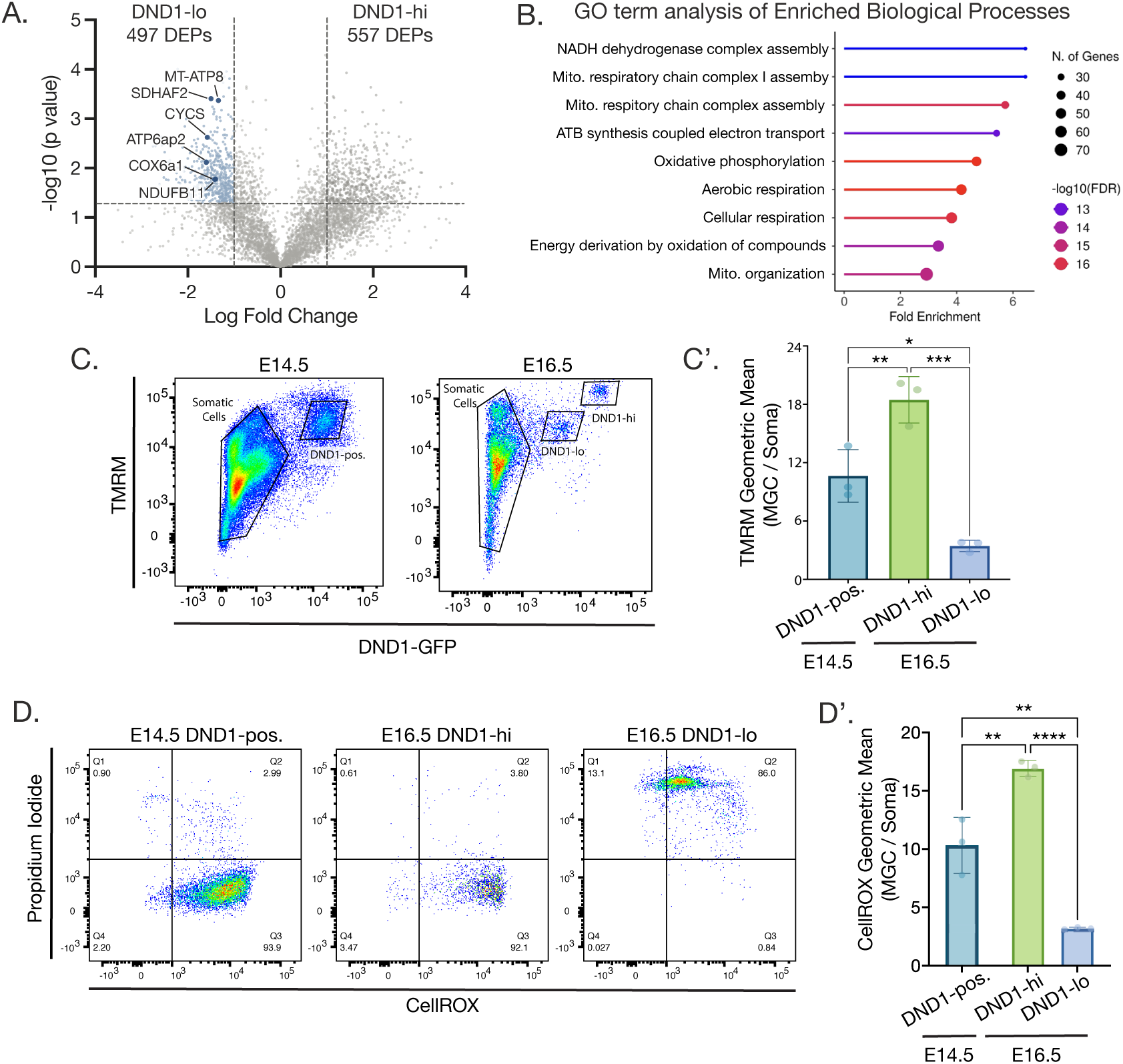
E16.5 DND1-lo cells show signs of mitochondrial dysfunction and decreased metabolism. ***(A)*** Volcano plot comparing the proteomes of DND1-hi versus DND1-lo cell populations at E16.5. Differentially expressed proteins (DEPs) are defined as having a -log10(p-value) > 1.3 and a Log2FC < 1. DND1-lo DEPs are highlighted in blue, and select mitochondrial proteins are labeled. ***(B)*** GO term analysis of DND1-lo DEPs. ***(C)*** Flow analysis of E14.5 and E16.5 gonadal cells stained with TMRM, a marker for positive membrane potential associated with mitochondrial health. *(C’)* Graph plotting the geometric mean of the MGC TMRM signal normalized to the TMRM signal of the gonadal soma. Significance based off ANOVA and Tukey’s multiple comparison’s test (\**P* < 0.05, \*\**P* < 0.01, \*\*\**P* < 0.0005, n = 3 litters). ***(D)*** Flow analysis of E14.5 DND1-positive, E16.5 DND1-hi, and E16.5 DND1-lo cells stained with propidium iodide and CellROX to detect ROS abundance. *(D’)* Graph plotting the geometric mean of the MGC CellROX signal normalized to the CellROX signal of the testis soma. Significance based off ANOVA and Tukey’s multiple comparison’s test (\**P* < 0.05, \*\**P* < 0.01, \*\*\*\**P* < 0.0001, n = 3 litters).

Because TMRM and MitoTracker dyes both depend on positive mitochondrial membrane potential to stain mitochondria, it was unclear from these assays whether the DND1-hi cells possessed more total mitochondria or if the mitochondria within he DND1-lo population have lost their membrane potential. Examination of mitochondria via immunofluorescent imaging at E16.5 showed differential TOMM20 signal between MGCs (Fig. S6C). Quantification of TOMM20 normalized fluorescent intensity correlated linearly with DND1-GFP. Notably, the linear correlation between DND1-GFP and TOMM20 was stronger (R^2^ = 0.91) than that of either marker and TRA98, suggesting the correlation between TOMM20 and DND1-GFP is not random (Fig, S6D, E). Taken together, these data suggest that DND1-lo cells possess fewer mitochondria compared to DND1-hi cells.

High levels of oxidative phosphorylation (OXPHOS) have been reported during G0 arrest, and disruption of G0 entry has been shown to cause a failure of OXPHOS upregulation and MGC apoptosis[2]. Thus, we aimed to examine metabolic differences between DND1-hi and -lo cells during G0 arrest. We stained cells with CellROX, a marker for ROS, and quantified ROS abundance using flow cytometry. Consistent with the idea that MGCs have increased OXPHOS during G0 arrest, we found that E14.5 and E16.5 MGCs had much higher levels of ROS production compared to the somatic cells of the testis (Fig. S6F, G). Within the MGC population, ROS levels appeared uniform at E14.5 (Fig. 5D). However, at E16.5, the DND1-hi cells had significantly higher levels of ROS compared to the DND1-lo cells, which were also positive for PI (Fig. 5D, D’), supporting the hypothesis that low metabolic levels during G0 arrest are associated with markers of cell death.

### Mitochondrial stress, but not DNA damage, is capable of prematurely inducing a DND1-lo MGC population

Thus far, we have established that the DND1-lo population experiences significant cellular damage by E16.5, including both an increase in mitochondrial dysfunction (Fig. 5D) and dsDNA breaks (Fig. 4D, E). However, whether one source of cellular damage is causal for formation of the DND1-lo population was still unclear. To answer this, we induced either mitochondrial or DNA damage at E14.5 (when we do not see DND1-hi and -lo populations) and measured changes in DND1 levels. When we “challenged” the E14.5 MGC mitochondria through the addition of the cytochrome C inhibitor, CCCP, the DND1-positive MGC population split into two foci (Fig 6A).

**Figure 6.**
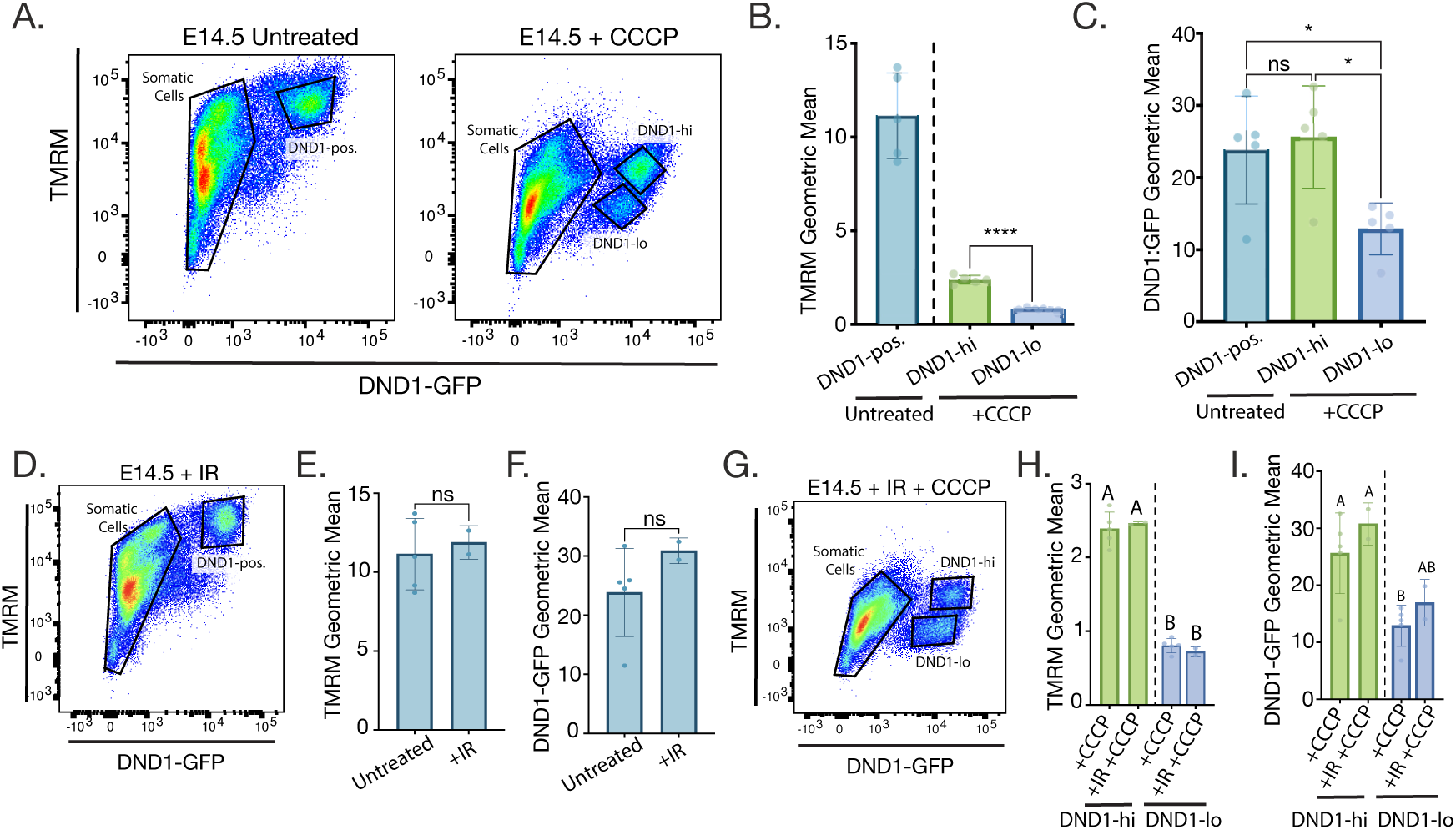
Mitochondrial stress, but not DNA damage, is capable of prematurely inducing a DND1-lo population at E14.5. ***(A)*** Flow analysis of untreated E14.5 MGCs or MGCs treated with a cytochrome C inhibitor (CCCP) used to challenge mitochondrial robustness. Cells stained with TMRM to measure mitochondrial membrane potential. ***(B)*** Graph plotting the geometric mean of the TMRM signal of E14.5 MGCs untreated or treated with CCCP. Unpaired t-test was performed to compare the two MGC foci that emerge after CCCP treatment (P < 0.05, n = 5). ***(C)*** Geometric mean of DND1-GFP signal between untreated E14.5 MGCs and the two MGC foci that appeared after CCCP treatment. The three groups were compared using ANOVA and Tukey’s multiple comparison’s test (\**P* < 0.05, n = 5 litters). ***(D)*** Flow analysis of testis cells from an embryo that had been treated with irradiation (IR, 200.0 cGy) to induce DNA damage. ***(E, F)*** Graph plotting the geometric mean of TMRM *(E)* or DND1-GFP *(F)* of E14.5 MGCs untreated or post-IR. Unpaired t-test was performed to compare the two MGC foci that emerge after CCCP treatment, (*P* > 0.05, untreated n = 5 litters, IR n = 2 litters). **(G)** Flow analysis of testis cells from an embryo that had been treated with irradiation (IR, 2 gray) and the cytochrome C inhibitor, CCCP. **(H, I)** Geometric mean of TMRM *(H)* or DND1-GFP *(I)* signal between MGCs treated with CCCP and MGCs treated with both CCCP and IR. The four groups were compared using two-way ANOVA, letters denote groups that are statistically similar (*P* > 0.05, untreated n = 5 litters, IR n = 2 litters). For all geometric mean quantification (*i.e., B, C, E, F, H, I*) the MGC geometric mean of TMRM or DND1-GFP was normalized to that of the gonadal soma.

One focus had a significant decrease in the geometric mean of the TMRM signal (Fig. 6B) and significantly lower DND1-GFP intensity, while the other focus was high for both DND1-GFP and TMRM signal (Fig. 6C). These data indicate that mitochondrial stress at E14.5 can prematurely induce formation of a DND1-lo MGC population.

We next tested whether induction of DNA damage could induce separate DND1-GFP populations by irradiating E14.5 pregnant females and performing flow analysis. While immunofluorescent staining for yH2AX confirmed the induction of dsDNA breaks in irradiated fetal testes (Fig. S7), flow analysis showed uniform expression of DND1-GFP within the MGC population (Fig. 6D).

Additionally, the geometric mean of TMRM and DND1-GFP in irradiated germ cells was statistically similar to that of the untreated controls (Fig. 6E, G), showing both TMRM and DND1-GFP levels are not responsive to DNA damage. However, if cells were treated with CCCP post-irradiation, two DND1-positive foci could once again be induced (Fig. 6G) with varying levels of TMRM (Fig. 6H) and DND1-GFP (Fig. 6I). Together, these data reveal that DNA damage is not capable of inducing a DND1-lo population, suggesting that mitochondrial stress is more likely an initiating source of cellular damage during G0 arrest.

## Discussion

Previously we reported that a proportion of MGCs becomes positive for the cell death marker, AnV, during G0 arrest[23]. However, the significance of this finding was unclear as no MGC apoptosis has been described during this period. Following up on a previous study reporting necrotic MGCs[17], we examined evidence for non-apoptotic cell death during G0 arrest. We found that MGCs display hallmarks of necroptosis around the time of birth (Fig. 2). To understand the basis for the appearance of these markers, we examined the health of the MGC population during late gestation and found accumulation of cellular damage beginning at E16.5 and corresponding with low expression of the RNA-binding protein, DND1 (Fig. 3). Proteomic analysis of the E16.5 DND1-hi and -lo MGC populations revealed an increase in DNA repair machinery in DND1-hi cells, including members of the MRN complex (Fig. 4). Meanwhile, the DND1-lo population harbored a significant increase in dsDNA breaks (Fig. 4) and exhibited characteristics of mitochondrial dysfunction (Fig. 5). Interestingly, induction of mitochondrial stress, but not DNA damage, was sufficient to prematurely induce a DND1-lo population at E14.5. These findings suggest that mitochondrial/metabolic dysfunction underlies the phenotype of DND1-lo cells.

Our work found the accumulation of cell-death associated markers within the DND1-lo MGC population. Previous work in the MGC field showed that differential protein expression during G0 arrest can lead to differences in MGC fate outcome. Seminal work from Law *et al*., demonstrated that high expression of the transcriptional repressor ID4 at E16.5 is a predictor of SSC fate[46]. ID4-hi cells harbor high levels of SSC-associated transcripts at E16.5. While we found that most of these genes were also higher within the DND1-hi population (Fig. S4A), this may be due to higher transcriptional levels in the DND1-hi cells. While rapid transcription can be damaging to the genome in some contexts[9], hypertranscription in PGCs has been associated with maintenance of PGC-specific gene expression and maintained genomic integrity[48, 49].

Interestingly, *Nanog* was one of the few cell-fate determining genes that was more abundant in the DND1-lo population. Previous work has shown that *Nanog* expression spikes at E14.5 and decreases by E16.5[50]. One possibility is that increased *Nanog* expression in the E16.5 DND1-lo cells is reflective of a developmental delay, possibly caused by declining cell viability. At the least, these findings reveal that fetal germ cells have high heterogeneity with respect to cellular health, a factor that should be considered in studies of germ cell programming.

Analysis of the DND1-hi and -lo populations during G0 arrest suggested a strong metabolic component to MGC survival. We found that G0-staged MGCs have higher levels of ROS than that of the testis soma (Fig. S6F, G). As ROS is a direct product of OXPHOS, this increase in ROS levels may reflect the metabolic shift to high OXPHOS known to occur during G0 arrest [2, 51]. Increased OXPHOS has been reported at several points in germ cell development including PGC and SSC differentiation, suggesting that germ cells experience metabolic shifts during times of transition and fate specification [51–54]. Furthermore, Du *et al.* found that errors in G0 entry led to a failure of OXPHOS upregulation, ectopic meiotic gene expression, and MGC apoptosis [2]. In the context of our work, this suggests that high OXPHOS during G0 arrest is important for MGC survival. We hypothesize that the DND1-lo population may arise from cells that are unable to sustain the metabolic demands of cellular reprogramming. These findings highlight the importance of metabolism in MGC reprogramming and survival during G0 arrest. Future efforts are needed to characterize the reason for differential metabolic states within the G0-staged MGC pool and to discover how OXPHOS functions to maintain MGC fitness and identity.

We observed that, between E14.5 and E16.5, MGCs that become DND1-hi show an increase in the abundance of healthy mitochondrial (Fig. 5C), which may account for their high ROS production. However, how and why the DND1-hi population increases their mitochondrial content between E14.5 and E16.5 while the mitochondria of the DND-lo population deteriorate remains unclear. One possibility is that differential MGC metabolic success stems from variable Sertoli cell support. Evidence suggests that Sertoli cells support the metabolic needs of germ cells. In *Drosophila,* germ cell survival is sustained by lactate produced by the supporting cyst cells and shuttled to the germ cells[55]. Work from our lab also showed that Sertoli cells supply lactate to PGCs prior to G0 entry (E11.5-E13.5)[56]. Thus, perhaps DND1-hi cells arise from MGCs that receive a higher nutrient supply from supporting cells. Rodriguez *et al.* concluded that MGC apoptosis during adolescence modulates MGC/Sertoli ratios in accord with the number of germ cells that can be supported by the Sertoli cell population[19]. We hypothesize that G0-staged MGC death may function similarly in that the number of surviving MGCs may be determined by the supportive capacity of the Sertoli cells. Further investigation is needed to understand the relationship between Sertoli cells and G0-staged MGC health.

We found evidence that neonatal MGCs display molecular characteristics of necroptosis (Fig. 2). Necroptotic cell death is best described in instances of infection or disease[31, 35, 37], and few physiological instances of necroptosis have been observed. While necroptosis has been implicated in SSCs removal in aged mice[57], it is surprising that a highly inflammatory cell death mechanism would be favored during development. One possible explanation is that, unlike apoptotic cell death, necroptosis does not require ATP, making it favorable in instances of ATP deprivation [58, 59]. Given the decreased mitochondrial health observed in the DND1-lo population, perhaps these cells are ATP deficient and thus more capable of necroptotic cell death than other cell death pathways. Additionally, we observed higher expression of *Cflar* within the DND1-lo population. Expression of mammalian *Cflar* can have complex effects as it encodes multiple gene products, but some studies have suggested that the *Cflar* encoded protein, cFLIP(s), can inhibit apoptosis and promote necroptosis[60, 61]. Interestingly, studies in *Drosophila* have shown pre-meiotic germ cells die through an unidentified non-apoptotic pathway[33, 34]. This raises the possibility that non-apoptotic cell death mechanisms such as necroptosis play a conserved role in germ cell development. It is unclear whether developmental necroptosis imparts any physiological benefit.

While G0-staged MGCs display hallmarks of necroptosis, there are many aspects of G0-staged MGC death that remain highly unusual. For instance, pMLKL is thought to be upstream of positive PI staining, yet we see accumulation of PI positive MGCs days prior to MLKL phosphorylation.

This raises the possibility that a distinct mechanism mediates MGC death during gestation that is still to be defined. We found that RIPK3’s RHIM domain is necessary for MLKL phosphorylation and nuclear HMGB1 loss (Fig. 2D, F), yet the protein acting upstream of RIPK3 is still unknown. Identifying the protein(s) that activate RIPK3 may be key to understanding the link between MGC death during late gestation and MLKL phosphorylation at the time of birth. Additionally, despite the progressive accumulation of DND1-lo cell damage across G0, we do not see any indicators of active cell removal until the time of birth, as shown through pMLKL (Fig. 2A). These data indicate that G0-staged MGC death occurs over a 4-day window (E16.5-P0), which raises numerous questions for further study. For instance, why G0-staged MGCs do not activate any cell death cascade prior to P0 remains unclear. One possibility is that the state of G0 arrest allows MGCs to overlook cellular damage until they prepare to re-enter the cell cycle. Staining for Ki67 showed that a portion of MGCs (12.58±7.81%) re-enter the cell cycle at P0 (Fig. S3B, C), coinciding with the time at which activation of a cell death pathway occurs. However, a more detailed analysis of G0 cell cycle re-entry is needed to definitively answer this question.

Finally, we could not capture a decrease in MGC number around birth. We hypothesize that this is because cycle re-entry occurs simultaneously. However, we cannot rule out the null hypothesis that DND1-lo MGCs recover from extensive cellular damage and survive. If this is the case, the accumulation of cell death-associated markers may represent a highly unique mechanism of MGC reprogramming. While we are not aware of any instances where cells survive accumulation of the markers we observed for cell death (*i.e.* PI-positive nuclear staining, phosphorylation of MLKL, and loss of nuclear HMGB1) there are examples of other cell types accruing substantial nuclear damage and surviving[62–64]. We hope that our present study lays the groundwork for future work to continue to explore how differential markers of MGC health act as significant regulators of MGC selection and programming.

## Materials and Methods

### Colony maintenance and timed matings

All mice were maintained in compliance with NIH guidelines, with approval from Duke University Medical Center IACUC (Protocol #A126-17-05). The *Dnd1^GFP^* line was generated by the Capel lab as previously described [22] on a B6SJLF1/J background (Jax strain# 100012) and outcrossed to CD1 mice (Charles River Labs strain code 022) approximately every six generations, then maintained homozygous (*Dnd1^GFP/GFP^*). The *Tnfr1/2* knockout mice were supplied by Dr. Edward Miao (Jax strain# 003243). *Ripk3 RHIM* knockout mice were supplied by Dr. Tomokazu Souma, who was kindly gifted these mice from Dr. Francis Chan [41]. *Zbp1* knockout mice were also supplied by Dr. Souma, who was kindly gifted these mice by Dr. Douglas Green [65]. *Rosa-mT/mG* mice were supplied by Dr. Terry Lechler (Jax strain# 007576). *Oct4-CreER mice* were purchased from Jax (Jax strain# 016829).For time mating, *Dnd1^GFP/GFP^* males were crossed to either *Dnd1^GFP/GFP^* females or CD1 females. Females were checked for plugs and staged as day E0.5 if positive. For neonatal mouse collections, breeding pairs were set up and checked for litters daily. The date of birth was recorded for litters and pups were collected.

### Isolation of DND1(+), DND1-hi, and DND1-lo cells via flow cytometry

For each assay, E14.5, E16.5, or E18.5 *Dnd1^GFP/GFP^* or *Dnd1^GFP/+^* testes were collected in fresh PBS. Testes were dissected away from the mesonephros and pooled (6-16 individual testes total depending on litter size). Limb tissue was collected in parallel to use as a GFP-negative control. All tissue was dissociated with Gibco TrypLE (Thermo Fisher Scientific, #12604013) and 0.5% Invitrogen Collagenase Type IV (Thermo Fisher Scientific, #17104019) at 37°C for 10 minutes.

After TrypLE treatment, tissue was resuspended in 500µL of PBS + 3% PSA (PBB). Tissue was gently pipetted up and down to create a single cell suspension, then passed through a 30μm cell strainer. Samples were analyzed using a BD LSRFortessa X-20 cell analyzer. Results were analyzed using FlowJo v10.10.0 Software (BD Life Sciences). Debris and doublets were excluded based on cell size. We identified DND1(+), DND1-hi, and DND1-lo cells using a defined gating strategy that has been previously published by our lab[23]. In brief, the Y-axis was set to SSC-A and the X-axis was set to DND1-GFP. Limb samples were used to define the cell scatter for GFP-negative populations, and cells appearing above 10^3^ on the DND1-GFP axis would be counted as DND1-GFP-positive, although there is some variation between samples.

### AnV and PI assay using flow analysis

E14.5, E16.5, or E18.5 *Dnd1^GFP/+^* testes were collected and dissociated as described in in the “*Isolation of DND1(+), DND1-hi, and DND1-lo cells via flow cytometry”* methods section with the following alterations: after TrypLE treatment, tissue was resuspended in 500μL of 1X Annexin-binding buffer (Annexin-binding buffer, 5X concentrate, Invitrogen, #V13246) and dissociated. Samples were then incubated with one drop of AnnexinV (AnV), Alexa Fluor 647 Ready Flow Reagent (Invitrogen, #R37175) and 1µL of 1.5mM propidium iodide (PI) (Thermo Fisher Scientific, #P1304MP) in the dark at 37°C for 15 minutes. Cells from the limb were also collected and prepared in parallel to use as unstained or single-stained controls. For AnV and PI single stained controls, limb cells were incubated with either AnV or PI in the dark at 55°C for 20 minutes. Following staining, all samples were passed through a 30μm cell strainer and analyzed using a BD LSRFortessa X-20 cell analyzer. The results were analyzed using FlowJo v10.10.0 Software (BD Life Sciences). Gating for DND1-hi and -lo populations was performed as described above. The gates for AnV and PI-positive cells were set based on the cell scatter from unstained limb and single-stained controls. Applying this gating to DND1-hi and -lo populations allowed us to collect cell counts within these quadrants which were used to for subsequent calculations.

### Imaging of live testis cords stained with DAPI

Testes were collected from E16.5 and P0 *Dnd1^GFP/+^* animals. The mesonephros and tunica were carefully removed, and testes were gently pipetted using a P1000 in PBS to “detangle” the testis cords. Incubation conditions for DAPI staining were developed based on published protocols for live-tissue staining as well as protocols for use of DAPI as a live/dead stain[27, 28]. Testes were incubated in Eppendorf tubes loosely covered with aluminum foil at 37°C in PBS+3% BSA (PBB) with 1µg/mL DAPI for 90 minutes. After incubation, tissue was washed 3×8 minutes in PBB while rocking in the dark at room temperature. After the final wash, excess media was removed and 20µL of Matrigel (Corning #356231) was pipetted on top if each testis. Matrigel was allowed to solidify for 20 minutes at 37°C, then covered with PBB. Immunofluorescence images were taken with a Zeiss LSM780 confocal microscope system using the associated Zen software. Image processing was performed using FIJI version 2.16.0/1.54p. The proportion of DAPI positive MGCs was calculated by counting the number of DAPI positive MGCs within one image frame for a given testis cord and dividing by the total number of visible MGCs.

### Proteomic analysis of E16.5 DND1-hi and -lo cells

Testes were collected from *Dnd1^GFP/GFP^* embryos at E16.5 and DND1-hi and -lo populations were isolated as described in the “*Isolation of DND1(+), DND1-hi, and DND1-lo cells via flow cytometry”* section. After cell sorting, cells were spun down (500xg, 4°C, 10 minutes) and flash frozen using liquid nitrogen. Samples were sent to the Duke Proteomics and Metabolomics Core for preparation prior to LC-MS/MS analysis. The samples were thawed and adjusted to contain 0.1% n-Dodecyl-β-D-maltoside (DDM) by adding 1μL of 1% DDM in 500 mM TEAB. Each sample’s total volume was then reduced with 1μL of 100 mM DTT and incubated at 80°C for 20 minutes, followed by a brief centrifugation. Alkylation was performed by adding 1μL of 250 mM iodoacetamide for 30 minutes at room temperature, after which the reaction was quenched with 1μL of 100 mM DTT. The samples were then subjected to overnight digestion with 2μL of trypsin (25 ng/μL) at 37°C and shaking at 850 rpm. To acidify, 1.5μL of 10% TFA was added to bring the final concentration to 1% TFA. A study pool QC (SPQC) was prepared by combining 3μL from each sample, except for the sample labeled “DND1-GFP low replicate 3,” from which only 1μL was used. For each sample, 5μL was diluted in 20μL of 0.1% formic acid, along with 1.5μL of the “DND1-GFP low replicate 3” sample and loaded onto Evotips. The samples were washed twice with 20μL of 0.1% formic acid prior to analysis.Quantitative LC/MSMS was performed on each sample using Evosep One LC interfaced to a ThermoFisher Orbitrap Astral. Raw MS data was demultiplexed and processed in Spectronaut 19 (19.7.250203.62635 Biognosys). A spectral library was built using direct-DIA searches which used a Uniprot *Mus muculus* database downloaded on 08/04/22, and appended contaminant sequences using FragPipe. For DIA analysis, default extraction, calibration, identification, and protein inference settings were used. Data was filtered at a 1% precursor and protein group false discovery rate (q-value). All precursors that met a q-value were used and protein abundances were calculated using the MaxLFQ algorithm. Data was log2-transformed in GraphPad Prism and a t-test was performed with or without Benjamini Hochberg FDR correction.

### GO term analysis

Proteins that were determined to have significant differential expression in the DND1-hi or the DND1-lo population were utilized for gene ontology (GO) analysis using ShinyG0 (https://bioinformatics.sdstate.edu/go/). Analysis was run with the background set to the E16.5 MGC proteome as identified from LC/MS analysis (all proteins identified from the DND1-hi and DND1-lo proteomic analysis). Pathway database was set to GO biological process; FDR cutoff was 0.05. Top ten pathways identified were reported.

### Protein network visualization

Protein interaction networks were generated using Cytoscape[66], with nodes representing significantly enriched proteins form the DND1-hi population by LC-MS/MS. High-confidence known interactions (score ≥ 0.7) among these proteins were retrieved from the comprehensive STRING database (https://string-db.org; [67]) and visualized within the network. To identify protein communities within each proximity proteome, the Markov Cluster Algorithm (MCL) implemented in STRING was employed, complemented by gene ontology (GO) analysis using ShinyGO[68]. Additional adjustments were made based on established protein functions.

### Neutral Comet Assay

Testes were collected from *Dnd1^GFP/+^* embryos at E14.5, E16.5, and E18.5 and dissociated as described in the “*Isolation of DND1(+), DND1-hi, and DND1-lo cells via flow cytometry”* methods section. Cells were sorted into DND1-hi and DND1-lo populations using a B-C Astros cell sorter as described previously[22]. This process was done in parallel with cells from dissociated limb tissue; limb sample was either untreated (negative control) or incubated at 55°C for 3 minutes to induce DNA damage (positive control). After cell sorting, a neutral comet assay was performed according to the specifications of Enzo’s Comet SCGE assay kit (#ADI-900-166) with the following specifications: cells were combined with 75µL of prewarmed LMAgarose in a 1:10 v/v ratio. Notably, we collected <50,000 DND1-hi cells at E18.5; in this case, all cells collected were combined with the LMAgarose and plated onto the comet slide. When running slides in the gel apparatus, slides were aligned equidistant from the electrodes, and the power supply was set to 1 volt/cm measured electrode to electrode. We ran the electrophoresis at 30V for 18 minutes. After slide preparation, images were taken using an Axio epifluorescent microscope on the EGFP laser with the laser power set to 20% and the exposer set to 500ms. Z-stacks were taken using interval setting with a z-step size of 5µm, and at least 10 z-stacks were taken per sample. Images were analyzed using FIJI software using max-projection of the z-stacks. Comet length was measured form the beginning of the comet head to the end of the visible tail.

### Flow analysis to assess mitochondrial health

TMRM, Mitotracker, and CellROX assays were performed using flow analysis to assess various markers for mitochondrial health. Testes were collected from E14.5 and E16.5 *Dnd1^GFP/+^* embryos were collected and prepared for flow cytometry as described in the “*Isolation of DND1(+), DND1-hi, and DND1-lo cells via flow cytometry”* methods section. Tissue from the limbs was also collected and used either as an unstained or single-stained controls. Cells were treated according to the specifications for each dye, described below. All results were analyzed using FlowJo v10.10.0 Software (BD Life Sciences). Gating and compensation were determined using unstained limb, single-stained limb, and unstained *Dnd1^GFP/+^* testes.

#### TMRM

MitoProbe TMRM Kit from ThermoFisher (#M20036). Single cell suspensions made from either testes or limb were incubated with 20nM TMRM reagent solution at 37°C for 30 minutes. For CCCP condition: samples were incubated with 50µM CCCP at 37°C for 5 minutes, then TMRM reagent was added, and samples were incubated at 37°C for an additional 30 minutes. For IR condition: prior to sample collection, the pregnant female was irradiated with the help of the Duke Radiology Core using a Cesium 137 source of radiation and treated with 200.0 cGy. The pregnant female was then allowed to recover for 2 hours before beginning embryo collection.

#### Mitotracker

MitoTracker Orange CM-H_2_ TMRos Kit from Invitrogen Molecular Probes (#M7511). Single cell suspensions were incubated with 200nM Mitotracker reagent at 37°C for 30 minutes.

#### CellROX

CellROX deep red reagent from Molecular Probes (#C10422). Single cell suspensions were incubated with 5µM CellROX reagent and 3µM PI at 37°C for 30 minutes. For a PI-positive single-stained control, cells from the limb were incubated with PI alone at 55°C for 20 minutes.

### Preparation and immunostaining of testis tissue

Testes were collected from embryonic or neonatal *Dnd1^GFP/+^* animals and prepared for whole mount staining or cryosectioning according to standard protocols previously described by our lab[22]. Immunostaining was also performed according to our standard protocol[22]. Immunofluorescence images were taken with a Zeiss LSM780 confocal microscope system using the associated Zen software. Image processing was performed using FIJI version 2.16.0/1.54p. All antibody information is provided in the table below.

| Primary Antibodies |  |  |  |  |  |
| --- | --- | --- | --- | --- | --- |
| Name | Host | Dilution | Vendor | Catalog # | Lot # |
| Anti-pMLKL(Ser345) | Rabbit | 1:1000 | Cell Signaling | 37333 | 4 |
| Anti-TRA98 | Rat | 1:500 | Abcam | ab82527 | 1098726-6 |
| Anti-yH2AX | Rabbit | 1:500 | Cell Signaling | 9718 | 21 |
| anti-GFP | Chick | 1:1000 | Abcam | ab13970 | 1018753-34 |
| Anti-TOMM20 | Rabbit | 1:500 | Abcam | ab186735 | 1096328-6 |
| Anti-HMGB1 | Rabbit | 1:1000 | Abcam | ab18256 | GR153222-1 |
| Anti-Ki67 | Rabbit | 1:500 | Cell Signaling | D3B5 | 3 |
| **Secondary Antibodies** |  |  |  |  |  |
| **Name** | **Host** | **Dilution** | **Vendor** | **Catalog #** | **Lot #** |
| Alexa Fluor 647-conjugated anti-rat | Goat | 1:500 | Jackson ImmunoResearch | 112-605-167 | 99468 |
| Alexa Fluor 488-conjugated anti-chick | Donkey | 1:500 | Jackson ImmunoResearch | 703-545-155 | 175023 |
| Cy3-conjugated affinitiPure Anti-Rabbit | Donkey | 1:500 | Jackson ImmunoResearch | 711-165-152 | 175018 |
| Alexa Fluor546 anti-rabbit | Donkey | 1:500 | Invitrogen | A10040 | 675498 |

### Immunofluorescent image quantification

#### TOMM20

Z-stacks were collapsed into a single image using slice-sum function in ImageJ. Germ cells were traced and the mean grey value (MGV) of TOMM20, DND1-GFP, TRA98, and DAPI was recorded for each cell. Because the MGC population is not dividing and each MGC should have a constant amount of DNA, the MGV of each marker was normalized by multiplying it by a DAPI normalization constant, which was calculated by dividing the mean DAPI signal by each individual DAPI measurement. The resulting value was referred to as the normalized fluorescent intensity (nFI).

#### pMLKL

Z-stacks were collapsed into a single image using slice-sum function in ImageJ. Each testis cord within an image was traced and the average grey value for all the pixels within the testis cord area was calculated this value was referred to as the mean grey value (MGV). MGV = (Sum of all grey values in the region) / (Number of pixels in the region).

#### HMGB1

Z-stacks were collapsed into a single image using slice-sum function in ImageJ. For each cell, the raw integrated density was measured for the nucleus and the whole cell. The nuclear area and cellular areas were also recorded. The background fluorescence was measured for each image by selecting an area without any cells and recording the mean signal. For each nuclei measured, the mean background signal was multiplied either by the nuclear area, then subtracted from the nuclear raw integrated density (nuclear raw integrated density - (mean background x nuclear area)). The same was done using the cellular area to remove the background signal from the cellular raw integrated density value. For each cell, the total cellular raw integrated density (minus background) was divided by the nuclear raw integrated density (minus background) to get a ratio of the cellular HMGB1 signal to the nuclear HMGB1 signal.

#### Counting MGC number

Four testes were sectioned per stage; for each testis, three to four sections were quantified. Z-stacks through each section were taken on a Zeiss LSM780 confocal microscope at the 10X objective and whole sections were imaged using the tile-scan function. In FIJI, max projections were taken of each Z-stack, and the total number of TRA98-positive cells were counted. The number of MGCs for each section for a given testis were averaged, and the average MGC per section number was plotted for each testis analyzed.

#### Ki67-positive MGC quantification

Images of cryosectioned testis tissue were imaged at 20X magnification. Per image analyzed, total MGCs (defined as TRA98-positive) were counted. Then the number of TRA98- and Ki67-positive cells were counted per frame, and the percentage Ki67-positive MGCs were calculated per image analyzed.

## Author Contributions

K.S., T.H., and B.C. designed research; K.S. and T.H. performed research and analyzed data; K.S. and B.C. wrote paper. All authors contributed comments.

## Acknowledgments

We would like to thank members of the Capel lab for their thoughtful discussion during the development of this project, and particularly Megan Harward for her contributions to mouse husbandry. We would like to thank Dr. Chantelle Evans and Olivia Conway for their feedback and for contributing TOMM20 antibody and CellROX reagents. The HMGB1 antibody was provided by Dr. Carolyn Coyne. Neonatal mice with *Ripk3 RHIM* and *Zbp1* mutations were provided by Dr. Koki Abe and Dr. Hiroki Kitai from the Souma lab. Dr. Souma was generously gifted these mice by Dr. Francis Chan (*Ripk3 RHIM* mutant) and Douglas Green (*Zbp1* mutant), with permission from Ken Ishii. Neonatal *Tnfr1/2* knockout mice were provided by Lupeng Li and Heather N Larson from the Miao lab at Duke University. Finally, we are grateful to the Duke Light Microscope Core Facility (LMCF), Duke Flow Cytometry Core Facility, and Marlene Violette and Matt Foster from the Duke University School of Medicine Proteomics and Metabolomics Core for their technical assistance on the mass spectrometry experiments. The following NIH sources of funding were used R01-HD103064 to B.C., F31-HD113207 to K.S., F32-HD113220 to T.H., R01-DK133369 to T.S, R01-AI175078, R01-AI181815, and R01-AI136920 to E.M.

## Supplemental Figures and Tables

**Figure S1.**
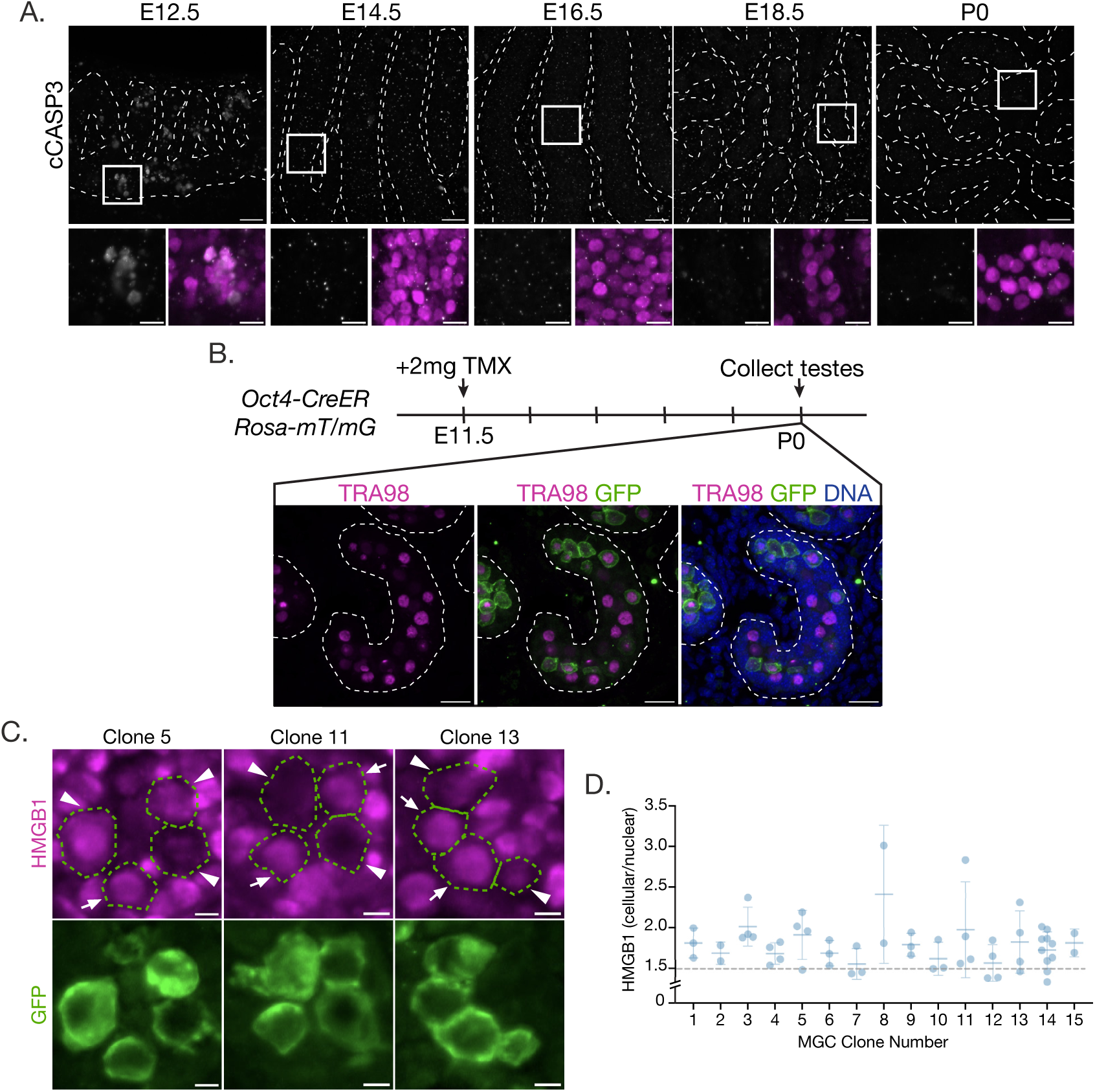
Neonatal MGCs show signs of a non-apoptotic cell death mechanism, independent of clonal origin. ***(A)*** Representative images of cCASP3 levels in whole mount testis cords between E12.5 and P0. Panels display cCASP3 (grey) and TRA98 (magenta). Testis cords outlined with white dashed lines. White boxes correspond to breakout images below. Scale bar for main images = 50µm, scale bar for breakout images = 20µm. ***(B)*** Schematic of experimental design for clonal MGC labeling. Images represent cryosections of *mT/mG* clonally labeled testes from P0 mice stained for GFP (green), TRA98 (Magenta), and DAPI (blue). Testis cords are outlined with dashed line, scale bar = 10µm. ***(C)*** Examples of MGC clones labeled with *mT/mG* reporter (GFP, green) and stained with HMGB1 (magenta). Clones outlined with dashed line in top panels. Arrows indicate healthy MGCs with nuclear HMGB1 signal; triangles point to MGCs with a loss of nuclear HMGB1. ***(D)*** Graph plotting cellular/nuclear HMGB1 ratio per clone. Data points represent individual cells within a clone. Error bars represent standard deviation.

**Figure S2.**
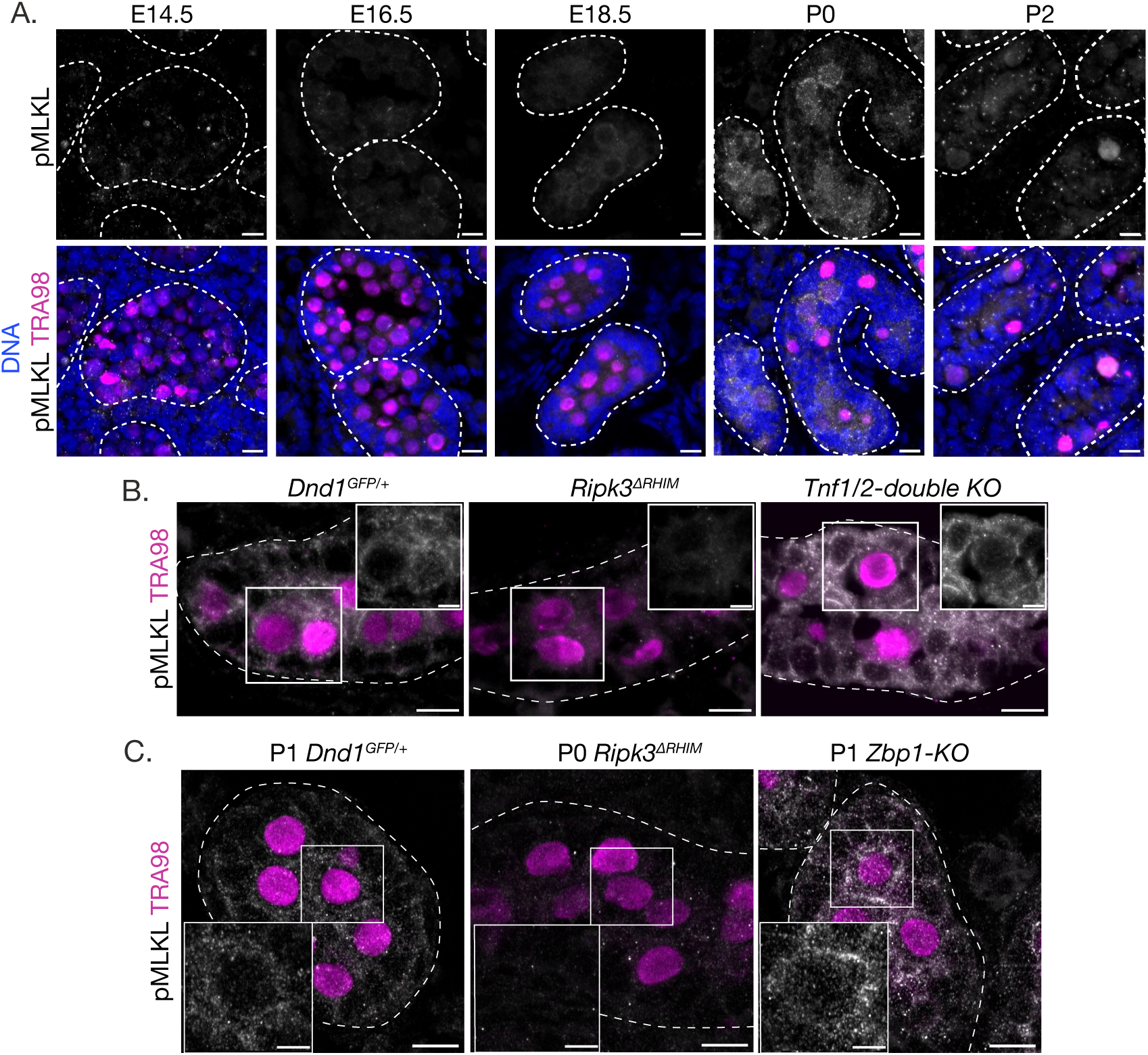
During G0 arrest, MGCs show signs of necroptotic activation that depends on RIPK3 RHIM function but not on TNF signaling or Z-DNA sensing via ZBP1. ***(A)*** Representative images of pMLKL levels across G0 arrest. ***(B)*** Images of P0 testis tissue from *Dnd1^GFP^*, *Ripk3^RHIMΔ^*, and *Tnfr1/2* knockout animals. *Tnfr1/2* knockout blocks TNF signaling. ***(C)*** Images of *Dnd1^GFP^*, *Ripk3^RHIMΔ^*, and *Zbp1* knockout animals. *Zbp1* knockout blocks Z-DNA sensing via ZBP1. Tissue was collected at either P1 (*Dnd^GFP^* and *Zbp1*-KO) or P0 (*Ripk3^RHIMΔ^*). All images display pMLKL (grey) and TRA98 (magenta) and represent testis cords outlined with dashed lines. Scale bars = 10µm, scale bars for higher magnification images = 5µm.

**Figure S3.**
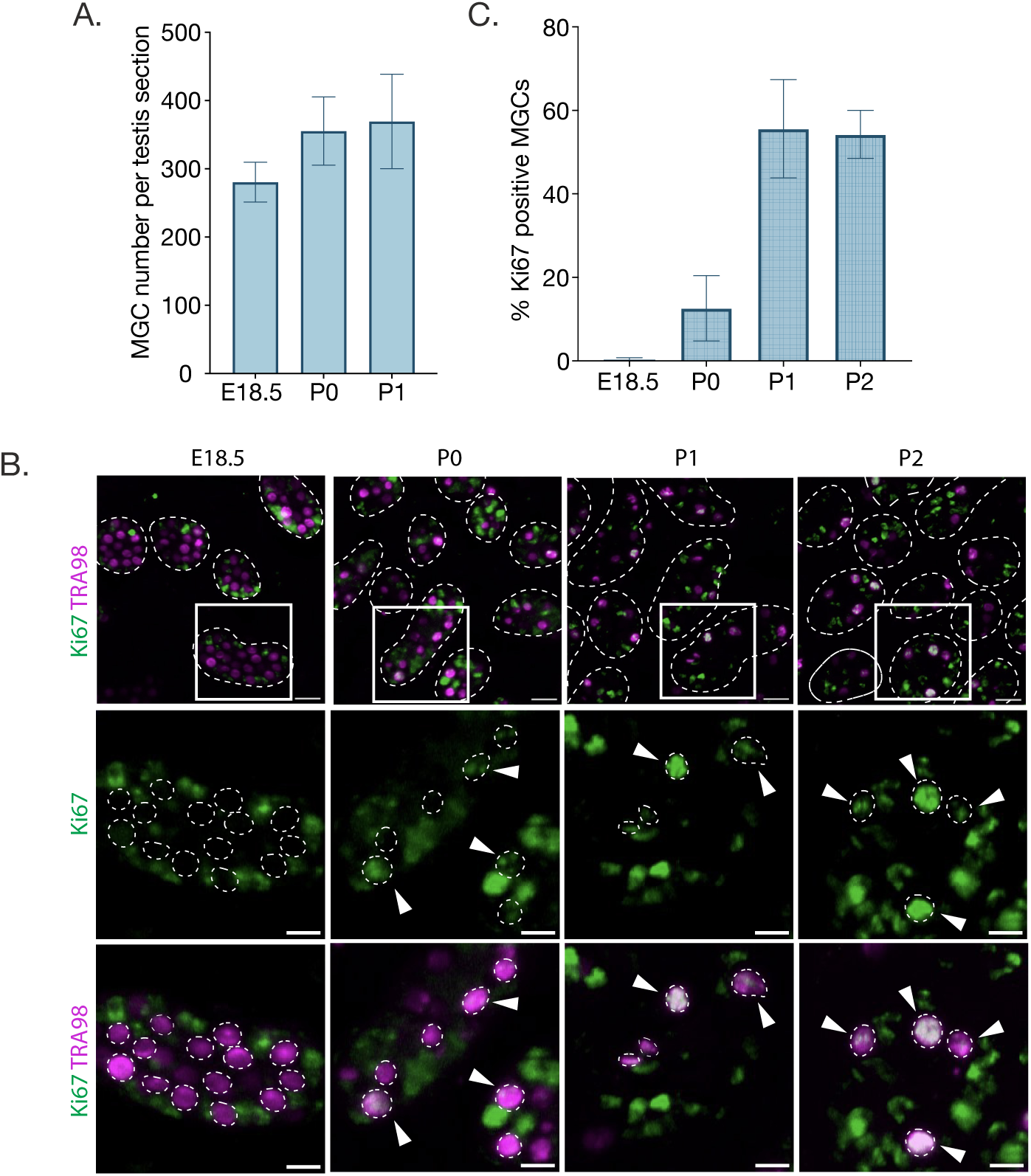
MGC numbers remain constant in the neonatal period, which may be due to cell cycle re-entry. ***(A)*** Bar graphs of average number of TRA98-positve cells per 10µm testis section (see methods). Counting was performed on multiple sections from 3-4 testes per stage, error bars represent standard deviation. ***(B)*** Representative images of Ki67 (green) and TRA98 (magenta) staining in cryosectioned tissue from 18.5, P0, P1, and P2 testes. Top panel shows testis cords outlined in dashed lines, scale bar = 20µm. Bottom two panels show magnified images at the indicated region, germ cells outlined with dashed line, arrows indicate Ki67-positive MGCs. Scale bar = 10µm. ***(C)*** Bar graph showing average percentage Ki67-positive MGCs per stage. Bars represent the average percentage of Ki67-positive MGCs per image analyzed (E18.5 n = 3, P0 n = 4, P1 n = 2, P2 n = 4 images per stage), error bars represent standard deviation.

**Figure S4.**
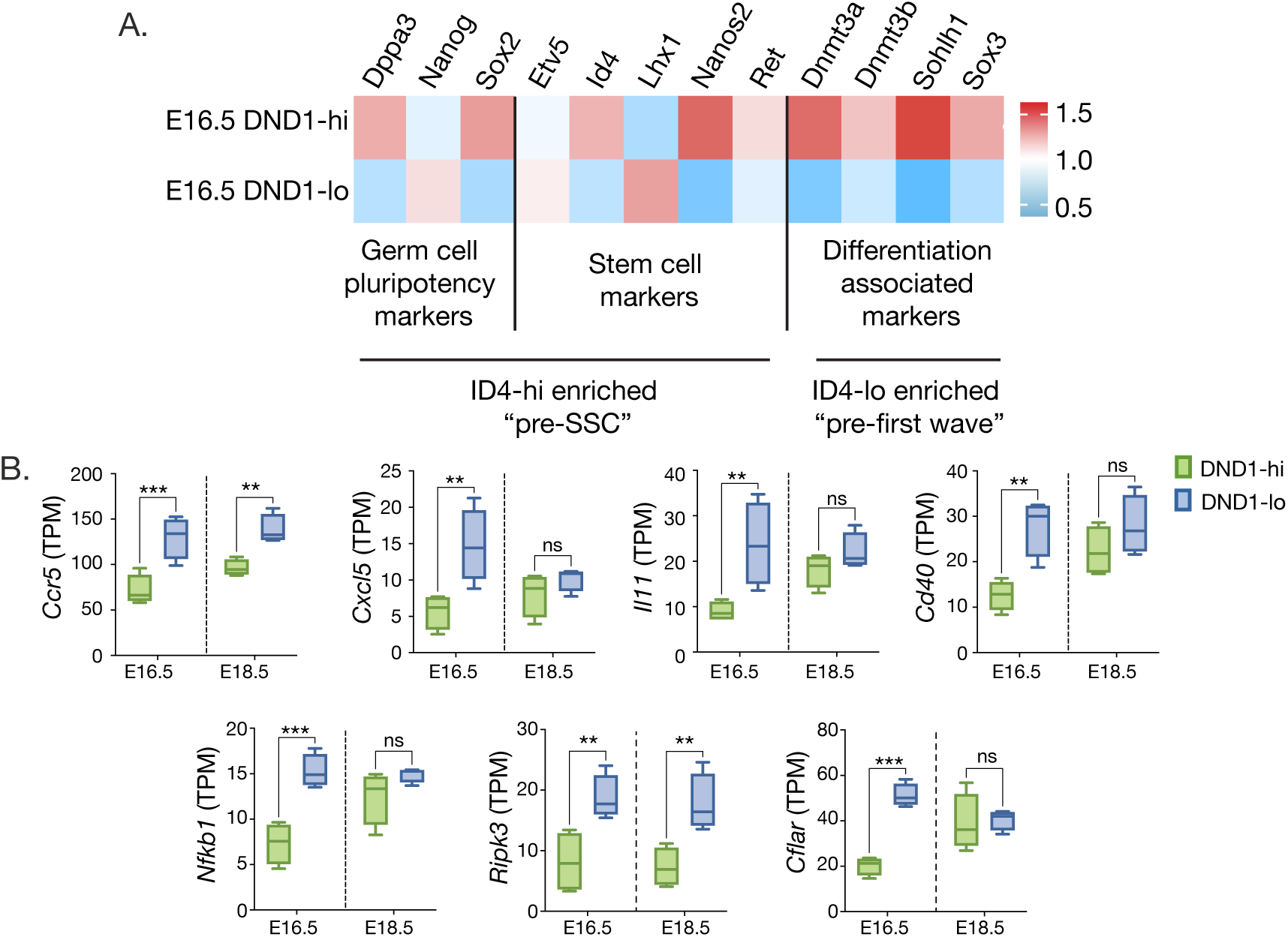
DND1-hi and -lo populations are not defined by “pre-SSC” or “pre-first wave” transcriptional profiles. ***(A)*** Heatmap displaying the relative expression of “pre-SSC” or “pre-first wave” genes between DND1-hi and DND1-lo populations at E16.5. Bulk RNA sequencing data was reanalyzed from Ruthig et al., [23]. Genes of interest were defined as “pre-SSC” or “pre-first wave” based on the definitions published by Law *et al.,* [46]. ***(B)*** Expression levels of various chemokine signaling and cell death related genes in DND1-hi and -lo cells at E16.5 and E18.5. Expression data taken from Ruthig *et al.*[23], significance based on 2-way ANOVA, \**P* < 0.05, *P* < 0.0.01, \*\*\**P* < 0.0001.

**Figure S5.**
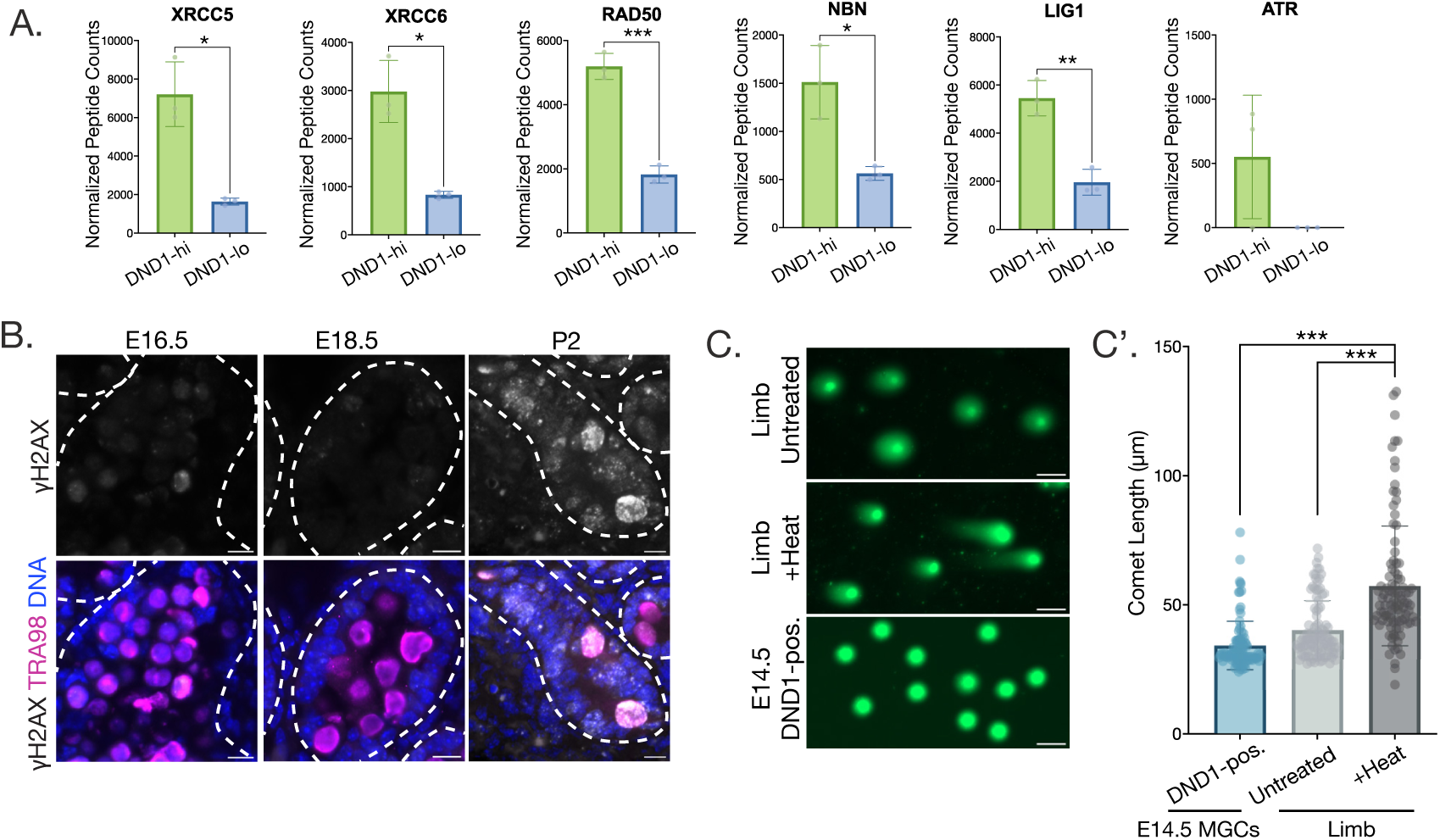
DND1-hi cells express higher levels of DNA repair proteins. ***(A)*** Bar graphs displaying the abundance of DNA repair proteins in DND1-hi and -lo cells at E16.5. Y-axis represents normalized peptide counts based on the findings of the DND1-hi -lo mass spectrometry analysis. Significance based on unpaired t test (*P < 0.05, **P < 0.005, ***P < 0.001). ***(B)*** Representative images of *γ*H2AX levels at E16.5, E18.5, and P2. Images taken of sectioned testis tissue, testis cords outlined with dashed white lines; bottom panels include DAPI and TRA98 staining. Scale bar = 10µm. ***(C)*** Representative images of comet assay performed on untreated limb cells (negative control), limb cells treated with heat (positive control), and flow sorted DND1-positive MGCs from E14.5 testes. Scale bar = 50µm*. (C’)* Quantification of comet tail length (µm) of DND1-positive E14.5 MGCs, untreated limb, and limb treated with heat. Significance based on unpaired t test, P < 0.0001 (n = 100 nuclei).

**Figure S6.**
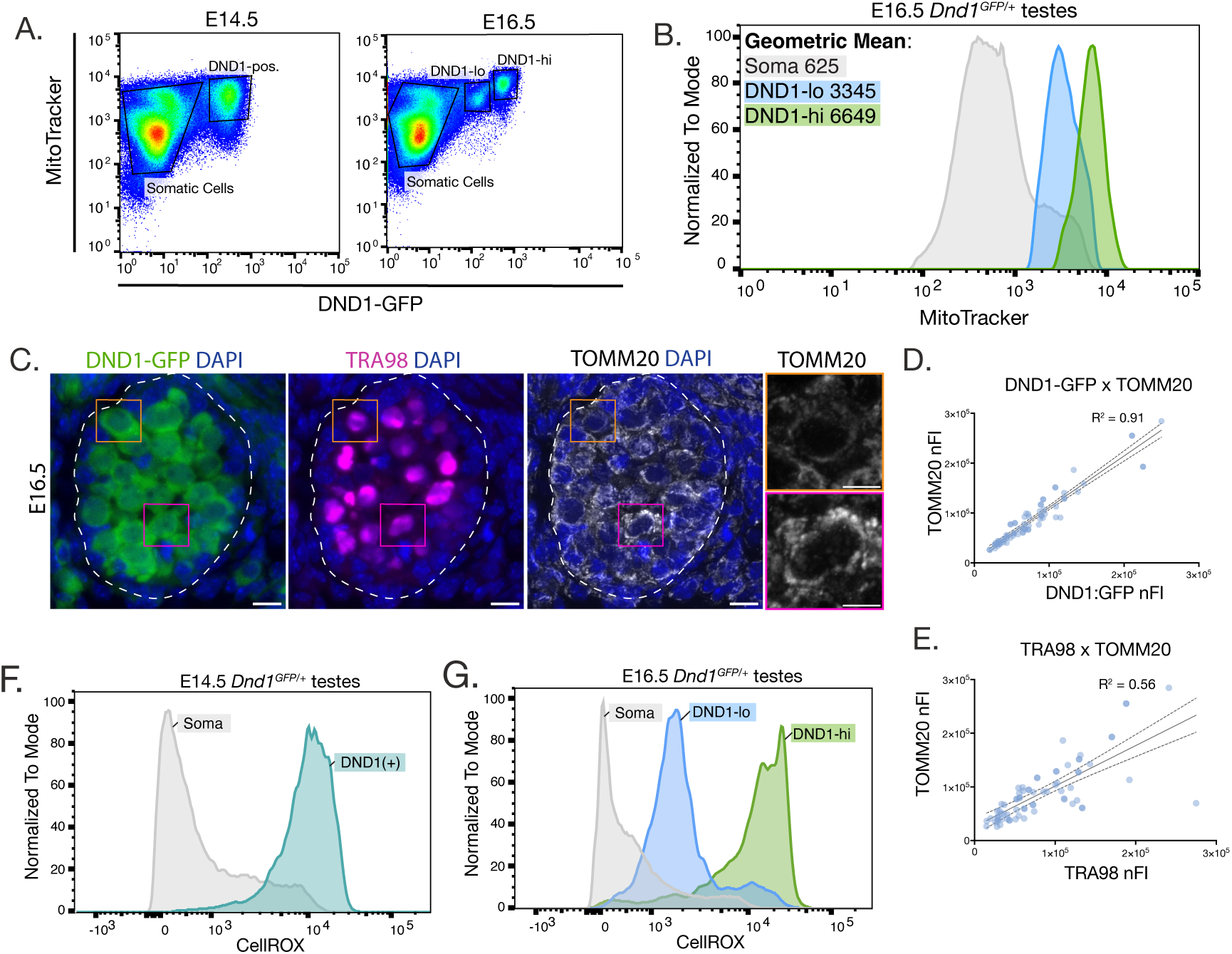
DND1-hi cells have more functional mitochondria and high levels of ROS. ***(A)*** Flow analysis of cells from E14.5 and E16.5 Dnd1^GFP^ testes stained with MitoTracker. ***(B)*** Geometric mean MitoTracker signal of somatic cell (grey), DND1-lo (blue), and DND1-hi (green) taken from E16.5 testes. ***(C)*** Representative image of sectioned testis cord displaying endogenous DND1-GFP signal and stained for TOMM20 and TRA98. Testis cord outlined with dashed white line. Scale bar of main image = 10µm. Scale bar of zoomed images = 5µm. ***(D-E)*** Graphs plotting normalized fluorescent intensity (nFI) of TOMM20 against DND1-GFP *(D)* and TOMM20 against TRA98 *(E).* All graphs were fit to a normal linear regression, R^2^ value represents the goodness of fit, dashed lines represent the 95% confidence interval. ***(F)*** Geometric mean of CellROX signal of somatic cells (grey) and MGCs (blue) at E14.5. ***(G)*** Geometric mean of CellROX signal of somatic cells (grey), DND1-lo cells (blue), and DND1-hi cells (green) at E16.5.

**Figure S7.**
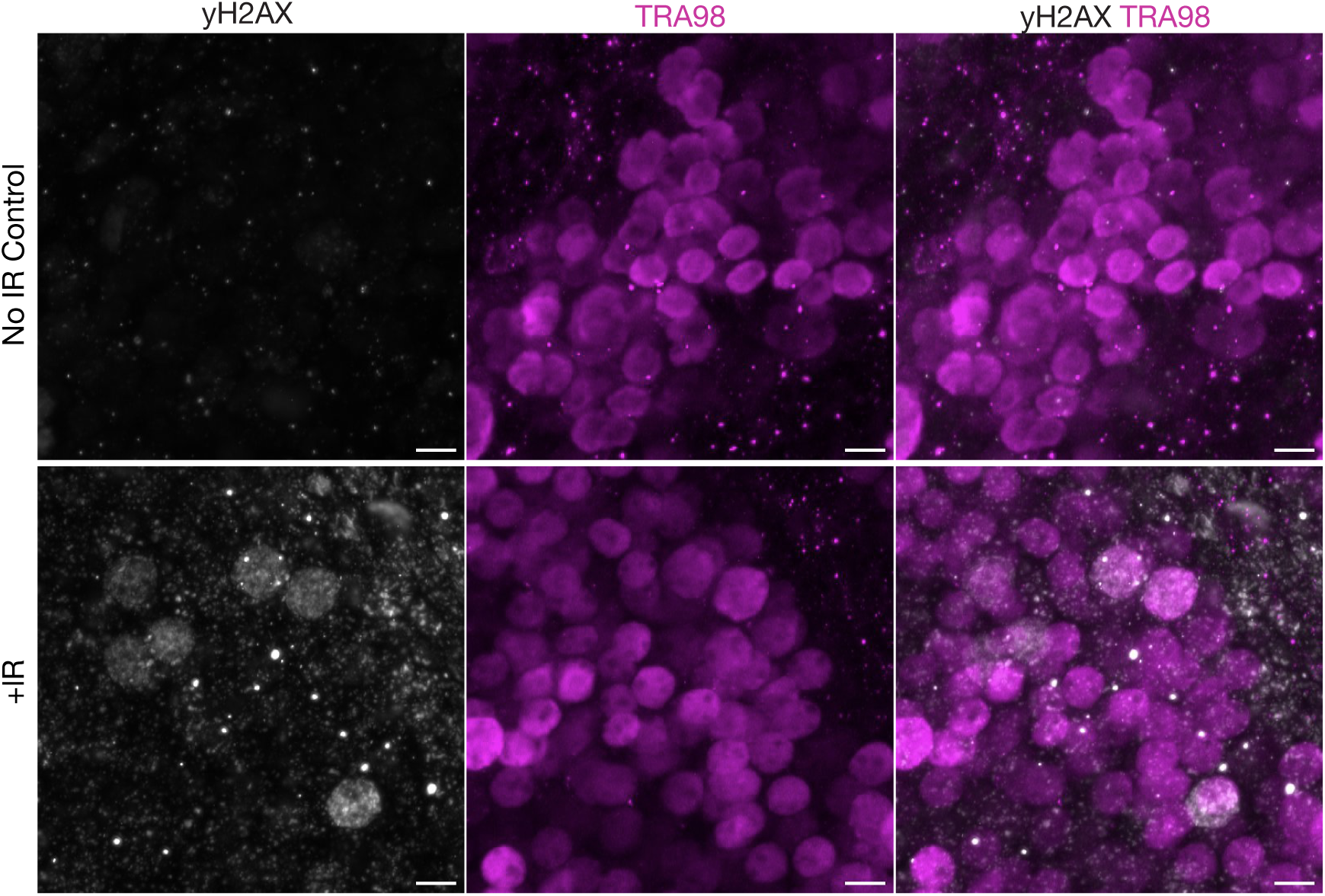
IR induces dsDNA breaks marked by yH2AX. **(A)** Representative images of yH2AX levels in whole mount testis cords from untreated E14.5 embryos and E14.5 embryos treated with 200.0 cGy irradiation. Panels display yH2AX (grey) and TRA98 (magenta). Scale =20µm.

**Table S1. Proteomic analysis of DND1-hi and -lo populations at E16.5.** Excel sheet with quantified protein abundance of DND1-hi and DND1-lo populations taken from mass spectrometry analysis. Table provides normalized counts for each replicate as well as statistical analysis of the differentially expressed proteins between the DND1-hi and -lo populations.

## References.

1. Seisenberger, S., et al., The dynamics of genome-wide DNA methylation reprogramming in mouse primordial germ cells. Mol Cell, 2012. 48(6): p. 849–62.

2. Du, G., et al., Proper timing of a quiescence period in precursor prospermatogonia is required for stem cell pool establishment in the male germline. Development, 2021. 148(9).

3. Dura, M., et al., DNMT3A-dependent DNA methylation is required for spermatogonial stem cells to commit to spermatogenesis. Nat Genet, 2022. 54(4): p. 469–480.

4. Goodwin, K., et al., Primordial germ cells experience increasing physical confinement and DNA damage during migration in the mouse embryo. bioRxiv, 2025.

5. Liu, S., et al., Setdb1 is required for germline development and silencing of H3K9me3-marked endogenous retroviruses in primordial germ cells. Genes Dev, 2014. 28(18): p. 2041–55.

6. Wang, D., et al., Active DNA demethylation promotes cell fate specification and the DNA damage response. Science, 2022. 378(6623): p. 983–989.

7. Rydberg, B. and T. Lindahl, Nonenzymatic methylation of DNA by the intracellular methyl group donor S-adenosyl-L-methionine is a potentially mutagenic reaction. EMBO J, 1982. 1(2): p. 211–6.

8. Stork, C.T., et al., Co-transcriptional R-loops are the main cause of estrogen-induced DNA damage. Elife, 2016. 5.

9. Palmer, B., et al., High-frequency transcription leads to rapid R-loop formation. J Biol Chem, 2025. 301(6): p. 108514.

10. Murphey, P., et al., Enhanced genetic integrity in mouse germ cells. Biol Reprod, 2013. 88(1): p. 6.

11. Walter, C.A., et al., Mutation frequency declines during spermatogenesis in young mice but increases in old mice. Proc Natl Acad Sci U S A, 1998. 95(17): p. 10015–9.

12. Milholland, B., et al., Differences between germline and somatic mutation rates in humans and mice. Nat Commun, 2017. 8: p. 15183.

13. Moore, L., et al., The mutational landscape of human somatic and germline cells. Nature, 2021. 597(7876): p. 381–386.

14. Kluin, P.M. and D.G. de Rooij, A comparison between the morphology and cell kinetics of gonocytes and adult type undifferentiated spermatogonia in the mouse. Int J Androl, 1981. 4(4): p. 475–93.

15. Pillai, R.S. and S. Chuma, piRNAs and their involvement in male germline development in mice. Dev Growth Differ, 2012. 54(1): p. 78–92.

16. Carmell, M.A., et al., MIWI2 is essential for spermatogenesis and repression of transposons in the mouse male germline. Dev Cell, 2007. 12(4): p. 503–14.

17. Wang, R.A., P.K. Nakane, and T. Koji, Autonomous cell death of mouse male germ cells during fetal and postnatal period. Biol Reprod, 1998. 58(5): p. 1250–6.

18. Nguyen, D.H., et al., Apoptosis in the fetal testis eliminates developmentally defective germ cell clones. Nat Cell Biol, 2020. 22(12): p. 1423–1435.

19. Rodriguez, I., et al., An early and massive wave of germinal cell apoptosis is required for the development of functional spermatogenesis. EMBO J, 1997. 16(9): p. 2262–70.

20. Rucker, E.B., 3rd, et al., Bcl-x and Bax regulate mouse primordial germ cell survival and apoptosis during embryogenesis. Mol Endocrinol, 2000. 14(7): p. 1038–52.

21. Russell, L.D., et al., Bax-dependent spermatogonia apoptosis is required for testicular development and spermatogenesis. Biol Reprod, 2002. 66(4): p. 950–8.

22. Ruthig, V.A., et al., A transgenic DND1GFP fusion allele reports in vivo expression and RNA-binding targets in undifferentiated mouse germ cellsdagger. Biol Reprod, 2021. 104(4): p. 861–874.

23. Ruthig, V.A., et al., The RNA binding protein DND1 is elevated in a subpopulation of pro-spermatogonia and targets chromatin modifiers and translational machinery during late gestation. PLoS Genet, 2023. 19(3): p. e1010656.

24. Balasubramanian, K., B. Mirnikjoo, and A.J. Schroit, Regulated externalization of phosphatidylserine at the cell surface: implications for apoptosis. J Biol Chem, 2007. 282(25): p. 18357–18364.

25. Span, L.F., et al., The dynamic process of apoptosis analyzed by flow cytometry using Annexin-V/propidium iodide and a modified in situ end labeling technique. Cytometry, 2002. 47(1): p. 24–31.

26. Vermes, I., et al., A novel assay for apoptosis. Flow cytometric detection of phosphatidylserine expression on early apoptotic cells using fluorescein labelled Annexin V. J Immunol Methods, 1995. 184(1): p. 39–51.

27. Sauvat, A., et al., Quantification of cellular viability by automated microscopy and flow cytometry. Oncotarget, 2015. 6(11): p. 9467–75.

28. Wallberg, F., T. Tenev, and P. Meier, Analysis of Apoptosis and Necroptosis by Fluorescence-Activated Cell Sorting. Cold Spring Harb Protoc, 2016. 2016(4): p. pdb prot087387.

29. Scaffidi, P., T. Misteli, and M.E. Bianchi, Release of chromatin protein HMGB1 by necrotic cells triggers inflammation. Nature, 2002. 418(6894): p. 191–5.

30. Volchuk, A., et al., Indirect regulation of HMGB1 release by gasdermin D. Nat Commun, 2020. 11(1): p. 4561.

31. Kim, D.Y., et al., RIPK1 Regulates Microglial Activation in Lipopolysaccharide-Induced Neuroinflammation and MPTP-Induced Parkinson’s Disease Mouse Models. Cells, 2023. 12(3).

32. Lei, L. and A.C. Spradling, Mouse primordial germ cells produce cysts that partially fragment prior to meiosis. Development, 2013. 140(10): p. 2075–81.

33. Yacobi-Sharon, K., Y. Namdar, and E. Arama, Alternative germ cell death pathway in Drosophila involves HtrA2/Omi, lysosomes, and a caspase-9 counterpart. Dev Cell, 2013. 25(1): p. 29–42.

34. Lu, K.L. and Y.M. Yamashita, Germ cell connectivity enhances cell death in response to DNA damage in the Drosophila testis. Elife, 2017. 6.

35. Chen, D., et al., RIP3-dependent necroptosis contributes to the pathogenesis of chronic obstructive pulmonary disease. JCI Insight, 2021. 6(12).

36. Lian, N., et al., Necroptosis-mediated HMGB1 secretion of keratinocytes as a key step for inflammation development in contact hypersensitivity. Cell Death Discov, 2022. 8(1): p. 451.

37. Huang, W., et al., Heat stress induces RIP1/RIP3-dependent necroptosis through the MAPK, NF-kappaB, and c-Jun signaling pathways in pulmonary vascular endothelial cells. Biochem Biophys Res Commun, 2020. 528(1): p. 206–212.

38. Ino, S., et al., Nuclear translocation of MLKL enhances necroptosis by a RIP1/RIP3-independent mechanism in H9c2 cardiomyoblasts. J Pharmacol Sci, 2023. 151(2): p. 134–143.

39. Yoon, S., et al., Necroptosis is preceded by nuclear translocation of the signaling proteins that induce it. Cell Death Differ, 2016. 23(2): p. 253–60.

40. Weber, K., et al., Nuclear RIPK3 and MLKL contribute to cytosolic necrosome formation and necroptosis. Commun Biol, 2018. 1: p. 6.

41. Moriwaki, K., et al., Distinct Kinase-Independent Role of RIPK3 in CD11c(+) Mononuclear Phagocytes in Cytokine-Induced Tissue Repair. Cell Rep, 2017. 18(10): p. 2441–2451.

42. Li, J., et al., The RIP1/RIP3 necrosome forms a functional amyloid signaling complex required for programmed necrosis. Cell, 2012. 150(2): p. 339–50.

43. Zhao, J., et al., Cell-fate transition and determination analysis of mouse male germ cells throughout development. Nat Commun, 2021. 12(1): p. 6839.

44. Pavlosky, A., et al., RIPK3-mediated necroptosis regulates cardiac allograft rejection. Am J Transplant, 2014. 14(8): p. 1778–90.

45. Baker, P.J. and P.J. O’Shaughnessy, Role of gonadotrophins in regulating numbers of Leydig and Sertoli cells during fetal and postnatal development in mice. Reproduction, 2001. 122(2): p. 227–34.

46. Law, N.C., M.J. Oatley, and J.M. Oatley, Developmental kinetics and transcriptome dynamics of stem cell specification in the spermatogenic lineage. Nat Commun, 2019. 10(1): p. 2787.

47. Al-Kafaji, G., M.A. Sabry, and M. Bakhiet, Increased expression of mitochondrial DNA-encoded genes in human renal mesangial cells in response to high glucose-induced reactive oxygen species. Mol Med Rep, 2016. 13(2): p. 1774–80.

48. Percharde, M., P. Wong, and M. Ramalho-Santos, Global Hypertranscription in the Mouse Embryonic Germline. Cell Rep, 2017. 19(10): p. 1987–1996.

49. Xia, B., et al., Widespread Transcriptional Scanning in the Testis Modulates Gene Evolution Rates. Cell, 2020. 180(2): p. 248–262 e21.

50. Yamaguchi, S., et al., Nanog expression in mouse germ cell development. Gene Expression Patterns, 2005. 5(5): p. 639–646.

51. Hayashi, Y., et al., Distinct requirements for energy metabolism in mouse primordial germ cells and their reprogramming to embryonic germ cells. Proc Natl Acad Sci U S A, 2017. 114(31): p. 8289–8294.

52. Lord, T. and B. Nixon, Metabolic Changes Accompanying Spermatogonial Stem Cell Differentiation. Dev Cell, 2020. 52(4): p. 399–411.

53. Sainz de la Maza, D., et al., Metabolic Reprogramming, Autophagy, and Reactive Oxygen Species Are Necessary for Primordial Germ Cell Reprogramming into Pluripotency. Oxid Med Cell Longev, 2017. 2017: p. 4745252.

54. Varuzhanyan, G., et al., Mitochondrial fusion is required for spermatogonial differentiation and meiosis. Elife, 2019. 8.

55. de la Maza, D.S., et al., Somatic cells compartmentalise their metabolism to sustain germ cell survival. bioRxiv, 2025.

56. Estermann, M.A., et al., Glycogen metabolism in mouse embryonic Sertoli cells sustains the germ line through the lactate shuttle. bioRxiv, 2025: p. 2025.07.23.666216.

57. Li, D., et al., RIPK1-RIPK3-MLKL-dependent necrosis promotes the aging of mouse male reproductive system. Elife, 2017. 6.

58. Liang, X., et al., Necroptosis, a novel form of caspase-independent cell death, contributes to renal epithelial cell damage in an ATP-depleted renal ischemia model. Mol Med Rep, 2014. 10(2): p. 719–24.

59. Ma, B., et al., ATP-Competitive MLKL Binders Have No Functional Impact on Necroptosis. PLoS One, 2016. 11(11): p. e0165983.

60. Sharp, D.A., D.A. Lawrence, and A. Ashkenazi, Selective knockdown of the long variant of cellular FLICE inhibitory protein augments death receptor-mediated caspase-8 activation and apoptosis. J Biol Chem, 2005. 280(19): p. 19401–9.

61. Ueffing, N., et al., Mutational analyses of c-FLIPR, the only murine short FLIP isoform, reveal requirements for DISC recruitment. Cell Death Differ, 2008. 15(4): p. 773–82.

62. Denais, C.M., et al., Nuclear envelope rupture and repair during cancer cell migration. Science, 2016. 352(6283): p. 353–8.

63. Häkkinen, H.-M., et al., In vivo nuclear envelope adaptation during cell migration across confining embryonic tissue environments. bioRxiv, 2025: p. 2025.06.05.658077.

64. Domingo-Muelas, A., et al., Human embryo live imaging reveals nuclear DNA shedding during blastocyst expansion and biopsy. Cell, 2023. 186(15): p. 3166–3181 e18.

65. Ishii, K.J., et al., TANK-binding kinase-1 delineates innate and adaptive immune responses to DNA vaccines. Nature, 2008. 451(7179): p. 725–9.

66. Shannon, P., et al., Cytoscape: a software environment for integrated models of biomolecular interaction networks. Genome Res, 2003. 13(11): p. 2498–504.

67. Szklarczyk, D., et al., The STRING database in 2025: protein networks with directionality of regulation. Nucleic Acids Res, 2025. 53(D1): p. D730–D737.

68. Ge, S.X., D. Jung, and R. Yao, ShinyGO: a graphical gene-set enrichment tool for animals and plants. Bioinformatics, 2020. 36(8): p. 2628–2629.

